# Glioblastoma–Neuroblastoma co-cultured multicellular spheroid model for evaluating Temozolomide response

**DOI:** 10.1101/2024.11.11.622957

**Authors:** Sivasubramanian Murugappan, Ajay Mali, Puja Sandbhor, Nanasaheb Thorat

## Abstract

Glioblastoma is the most lethal primary brain tumor, and its resistance to therapy is increasingly understood to arise not from tumor cells alone but also from their integration into the surrounding neural circuitry. However, no existing preclinical spheroid model isolates the direct effect of neuronal cells on glioblastoma’s drug response within a rapid, scalable 3D system. We developed a heterotypic 3D co-culture spheroid combining glioblastoma (U87-MG) with a neuronal lineage population (SH-SY5Y) via the agarose liquid overlay method to directly understand this gap. Subsequently, growth, morphology, hypoxia, and temozolomide (TMZ) response over 14 days against glioblastoma only and neuroblastoma only spheroids were characterized. The co-culture adopted a glioblastoma like compact, chemoresistant architecture yet, contrary to what this architecture alone would predict, it showed measurable chemosensitivity at every TMZ concentration tested, including 10µM, a dose at which neither monoculture responded at all. This reveals that the neuronal lineage component actively sensitizes glioblastoma to chemotherapy independent of spheroid architecture. Drug-response data from glioblastoma only models may therefore not reflect true tumor behavior within its native neuronal environment, underscoring the need for neuron inclusive platforms in preclinical glioblastoma drug screening.

**GRAPHICAL ABSTRACT.**
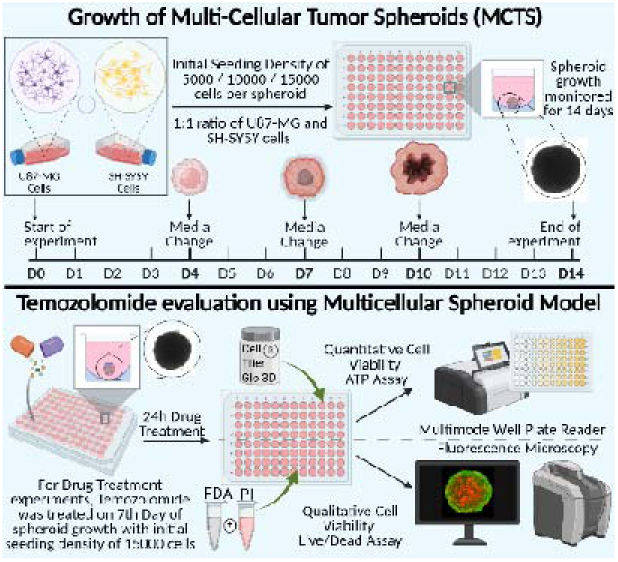
Schematic representation of the formation of spheroids and the evaluation of Temozolomide on it.

## 1. Introduction

Diffuse gliomas are primary brain tumors originating from glial or neuronal progenitor cells and are clinically challenging to treat [1]. Despite standard of care treatments including surgery, adjuvant temozolomide-based chemotherapy, and radiotherapy, gliomas typically showed poor prognosis due to extensive infiltration into the healthy brain parenchyma. Among gliomas, the most common type, glioblastoma (GBM), has an average survival of approximately one year, with 90% of patients experiencing relapse despite clinical care [2–4]. Moreover, seizures, epilepsy, and focal neurological deficits are major clinical manifestations seen in almost 30-60% of GBM patients either at diagnosis or across the disease progression, and contributing to neurological decline and neurotoxicity beyond the tumorigenesis effect [5].

A key reason this prognosis has not moved is that GBM does not behave as an isolated mass of malignant glial cells, it behaves as a tissue that has intervened itself into the neural circuitry around it within the central nervous system. In recent years, significant attention has been given to studying the interaction between GBM and neurons, to understand how tumor destroys surrounding neuronal tissue as collateral damage during the disease course. Evidence has shown that bidirectional crosstalk between the neurons and glioma cells promote neuronal circuits remodelling by inducing hyperexcitability, which in turn impairs the balance of neurotransmission, thereby promoting tumor survival and aggressive infiltration into surrounding brain [6,7]. Interestingly, several studies have shown that exacerbated neuronal hyperexcitability also contributed to pre-operative seizures and epilepsy episodes in GBM patients [8,9]. Additionally, clinical observations from site-specific tumor biopsy tissue has reported that GBM occur more frequently in brain regions with increased neuronal activity and correlating it with poor patient survival and performance on neurological scoring [10,11]. Furthermore, the glioma cells hijack neural tissue by extending neurite-like ultra-long membrane protrusions known as tumor microtubes, which in turn leads to early and rapid dissemination of tumor cells into surrounding brain [8,12,13]. Nonetheless, the cellular connection formed by these tumor microtubes is reported to be key mediator of resistance to chemotherapy in GBM [14]. In addition to tumor microtubes, the electrochemical synapses such as glutamatergic synaptic input mediated by α-amino-3-hydroxy-5-methyl-4-isoxazolepropionic acid receptors (AMPARs) promote invasion and proliferation of glioma cells via increased calcium entry through the tumor microtubes. In this regard, Cueto-Urena and collaborators have reported that blockade of this synaptic inputs suppress migration of glioma cells and induce their apoptosis [15]. However, despite the crucial role of GBM-neurons crosstalk, the evaluation of cellular functionality has not been performed from therapeutic perspective using acute tumor-neuron models, allowing glioma-derived cues to influence neurons and cell type specific effects on drug sensitization studies. Together, these findings underscore the importance of understanding glioma-neuron interactions and offering an insight into how glioma cells influence neurons during initial tumor infiltration which is key driver of disease progression, resistance to chemo drugs and recurrence.

To investigate neuronal response to GBM, the preclinical workflow has instead relied on two-dimensional (2D) co-cultures or tumor induced animal xenograft models [16]. However, both have well-known limits i.e. 2D culture attenuate the cell-cell/cell-matrix interactions that define tumour architecture and microenvironment while the animal models carry ethical and cost constraints. Moreover, high variability and species-specific differences for instance, human astrocytes alone differ substantially in size and function from rodent astrocytes [17,18] raise concerns regarding the clinical relevance of findings from preclinical outcomes. Furthermore, three-dimensional (3D) multicellular tumour spheroids (MCTS) address some of this challenges by bringing into the effect of essential cues like oxygen, nutrient diffusion, and drug-response that better recapitulate in vivo tumour physiology [19]. Although, organoid systems move forward, but require extended maturation period (several months), laborious, and offer negligible control over neuron-to-glioma ratios [20], limiting their utility in rapid throughput drug screening.

This highlights the specific gap that remains unresolved that is no existing spheroid model, model the acute stage of the direct effect of a neuronal-lineage population on GBM’s drug response, independent of tumor vasculature, immune infiltrate, or the month long/extended maturation that organoid systems require. Whether the existence of neuronal cells influence how GBM responds to chemotherapy rather than passively co-existing with it remained untested within a controlled, and scalable 3D system.

In present study, we address this gap directly. We developed a heterotypic 3D co-culture spheroid combining glioblastoma (U87-MG) with a neuronal-lineage population (SH-SY5Y) using the liquid overlay method, chosen specifically for its reproducibility and scalability over organoid culture [21]. Using this system, we asked two questions: first, does the presence of neuronal-lineage cells change GBM’s growth, architecture, and hypoxic profile in 3D, a baseline that matters because hypoxic, poorly perfused architecture is itself a known key driver of chemoresistance in GBM; and second, does it change GBM’s sensitivity to temozolomide (TMZ), the first-line chemotherapy agent for this condition. To answer this, we assessed the co-culture against GBM-only and neuroblastoma-only spheroids across 14 days of growth and a logarithmic range of TMZ concentrations.

As we show below, the neuronal-lineage component does not passively coexist with GBM in the spheroid it produces a measurable sensitization. Interestingly, the 3D spheroidal co-culture responded to TMZ at concentrations (10 µM) at which neither monoculture showed any response at all, a low-dose sensitivity that is invisible to every glioblastoma-only model currently used for drug screening. This outcome has a direct practical implication from the therapeutic perspective: drug efficacy data from GBM monocultures may not reflect how a tumour actually responds within its native neuronal environment. Closing this gap requires a neuron-inclusive, scalable 3D system established in this study. This platform provides a promising tool for modelling early-stage of glioma-neuron interaction preclinically allowing both therapeutic vulnerability and the resulting neurotoxicity to be tested.

## 2. Methodology

### 2.1 Cell culture and Formation of Spheroids

The glioblastoma and neuroblastoma cells were provided by Prof. James Curtin (Technological University Dublin) and Prof. Dennis Barry (Trinity College Dublin), respectively. The cells were cultured in a DMEM media supplemented with 1% penicillin-streptomycin, 1% L-Glutamine (Merck Life Sciences) and 10% Fetal Bovine Serum (FBS) (Cytiva HyClone). The Phosphate Buffer Saline (PBS) and Trypsin needed for the washing and detachment of cells were also obtained from Merck Life Sciences. The cells were maintained at 37°C and 5% CO_2_. Initially, Flat-bottomed 96 well plates (SPL Life Sciences) were coated with 1% agarose solution (Fisher Scientific). Then, cell suspensions with cell densities of 5000, 10000, and 15000 per well were prepared in 200 µL medium for U87-MG and SH-SY5Y. Equal amount (100 µL each) of U87-MG and SH-SY5Y cells were used to develop the co-cultured spheroid model. The spheroids were grown for 14 days. Media was changed on day 4, 7 and 10 after imaging. While changing media, only 100 µL of the old spent media was removed to not perturb the spheroids and dissociate them. Subsequently, fresh 100 µL media was added gently to the walls of the well to minimize the stress on the spheroids. Phase contrast microscope images of the spheroids were taken each day using a Keyence BZ-X810 Microscope.

### 2.2 Cell Track Assay

Both U87-MG and SH-SY5Y cells were tracked with the Cell Trace Far red and CFSE dyes, respectively. The protocol was followed as per manufacturer’s instructions. Briefly, after trypsinization and centrifugation, the supernatant media was removed from the cell pellet. Then, 1 mL PBS and dyes from stock solution was added such that the overall concentration was 10 µM. The cells were then incubated at 37°C for 20 minutes. Then 5 mL media was added to it, to remove excess dye. The supernatant was removed following centrifugation and was added with 1 mL media. Finally, total of 15000 cells/well in 1:1 ratio was seeded for spheroids.

### 2.3 Live/Dead Assay

Live/Dead Assay, a qualitative way of assessing the cell viability of spheroids, was performed on day 7, 10 and 14. Briefly, 150 µL of media was removed from each well, washed with PBS and were added with 10 µL of Propidium Iodide (PI) followed by Fluorescein Diacetate (FDA) dissolved in PBS and Acetone, respectively. The concentration of dyes was kept constant at 0.5 mg/mL. After an incubation period of 10 minutes, the spheroids were washed twice with PBS and taken directly for fluorescence imaging using a Keyence BZ-X810 Microscope.

### 2.4 Hypoxia – PI imaging

Briefly, 150 µL of medium was removed from each well, leaving 50 µL. Subsequently, 50 µL of Hypoxia Image-iT dye solution (prepared in culture medium) was added to each well, resulting in a final dye concentration of 5 µM in a total volume of 100 µL. The spheroids were incubated at 37 °C for 50 minutes. Thereafter, 10 µL of 0.5 mg/mL PI was added and the spheroids were incubated at 37 °C for a further 10 minutes. Finally, the spheroids were washed twice with PBS prior to imaging.

### 2.5 Drug Treatment

The spheroids were treated with Temoside® formulation obtained from Cipla Ltd, India at a logarithmic range to know the efficacy range of the drug in spheroid models developed. The concentration was based on the amount of active pharmaceutical ingredient – Temozolomide (TMZ). The final drug concentration was maintained at 10, 100, and 1000 µM.

#### 2.5.1 ATP Assay

On the 7th day of spheroid growth, 150 µL spent medium was removed from the 200 µL media in each well and 50 µL of drug dissolved media was added. After 24 h, the spheroids were added with 100 µL of Cell-Titer Glo 3D®, incubated at room temperature for 30 minutes. The luminescence values were obtained using BioTek Synergy H1 multimode plate reader.

#### 2.5.2 Live/Dead Assay

On the 7th day of spheroid growth, 100 µL spent medium was removed from the 200 µL media in each well and 100 µL of drug dissolved media was added. On Day 8, the spent media was removed and washed with PBS, incubated with PI and FDA and imaged.

### 2.6 Statistical Analysis

All image measurements were made using ImageJ. The circularity obtained from ImageJ followed the formula,

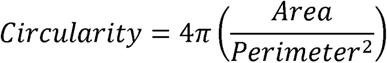

All statistical analyses were performed using either Python or GraphPad Prism software version 10.2.0. Repeated measures Two-way ANOVA statistic method with post-hoc Tukey test was used. P < 0.05 was considered significant (n = 3).

## 3. Results and Discussion

Tumor’s native neuronal environment plays a substantial role in development of preclinical glioblastoma (GBM) models to advance our understanding of how tumor progression along with their neuronal lineage during the disease course, and response to chemotherapy more analogous to the *in vivo* setting [22]. However, with the established GBM platforms, determining the most relevant model for assessing the effect of standard of care on tumor infiltration and therapy resistance is challenging specifically in neuronal context. For instance, conventional 2D monolayer system have laid the foundation of tumor biology with many key findings over the past decades [23]. Conversely, they rarely account the influence of neuronal environment in GBM behavior, despite growing evidence of structural and functional integration of GBM cells into surrounding neural circuitry through mechanisms including tumor microtubes, and synaptic inputs mediated through neuron-glioma crosstalk [8,11]. Furthermore, clinical manifestation of glioma associated neuronal excitability such as seizures, and epilepsy suggested the involvement of neuron-glioma interactions in growth and invasion of GBM [24]. Therefore, to directly assess whether this neuronal context influence GBM’s progression and drug response, the present study established a heterotypic 3D spheroid model by co-culturing glioblastoma (U87-MG) with a neuronal-lineage population (SH-SY5Y). Glioblastoma only and neuroblastoma only spheroids were grown in parallel as reference models using the liquid overlay method, and all three groups were characterized for growth kinetics, morphological assessment, spatial organization, and hypoxic profile over 14 days. This is crucial for establishing the architectural baseline necessary before evaluating chemosensitivity, since spheroid circularity and compaction, and hypoxia are themselves established drivers of drug penetration and resistance in 3D tumor models as supported by previous studies [25,26].

### 3.1. Spheroid growth and morphology

U87-MG spheroids maintained a compact, spherical architecture across all three seeding densities (Figure 1a), consistent with strong cell-cell adhesion driving early spheroid consolidation. At the liquid-air interface due to gravity, cell clusters were observed to form independently before migrating and fusing into the larger spheroids over period (Figure S1). This self-assembled behavior has been widely reported for glioma spheroid formation via the liquid overlay method. By day 14, the 5000, 10000 and 15000 cell groups reached diameters of approximately 400 µm, 500 µm and 600 µm respectively. All densities underwent an initial contractile phase over the first 3-4 days, during which projected area fell by 25-44 % depending on seeding density (Figure 4a) supporting that cell-cell adhesion, instead of proliferation, dominates the earliest stage of GBM spheroid formation. This finding was in-line with prior reports of an adhesion-driven compaction phase preceding proliferative expansion in tumor spheroids [27,28].

**Figure 1.**
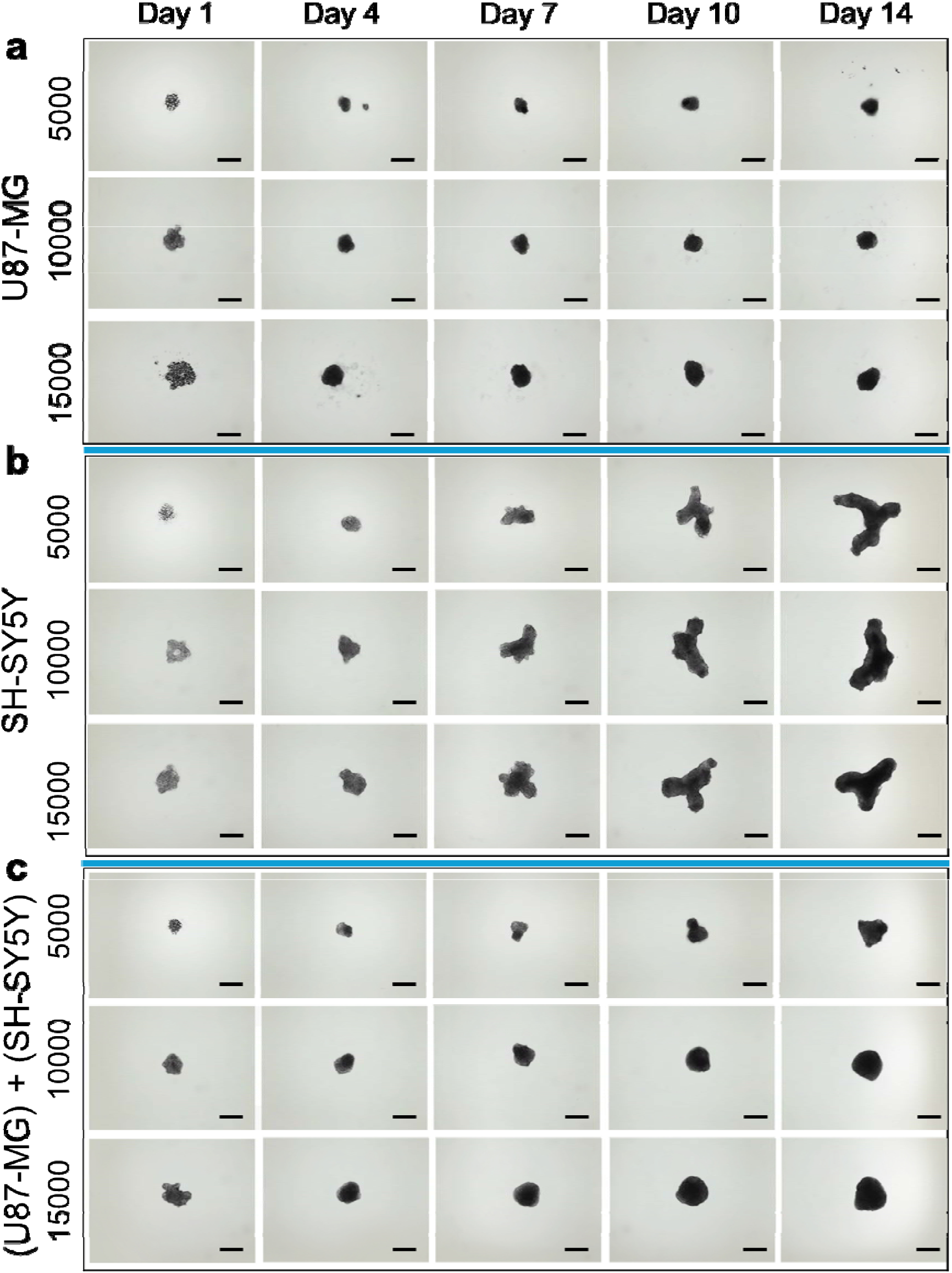
Spheroid growth over 14 days with cell seeding densities of 5000,10000 and 15000 cells: U87-MG (a), SH-SY5Y (b) and co-cultured spheroid (U87-MG)+(SH-SY5Y)) (c). Scale 500µm. (Images processed using Adobe Photoshop). All 14 day images in Figure S1.

In sharp contrast, SH-SY5Y (neuroblastoma) spheroids across all three densities adopted an irregular, elongated architecture rather than a compact sphere. SH-SY5Y cells were selected as a neuronal surrogate because they retain key neuronal characteristics while providing a practical and reproducible alternative to primary or iPSC derived neurons. Unlike primary neurons, SH-SY5Y cells proliferate readily and avoid variability associated with donor source, isolation procedures, and differentiation protocols. These advantages make them well suited for developing reproducible neuron-glioblastoma co-culture models for drug screening. We observed that this branching pattern closely resembled neurite outgrowth, consistent with the neuronal lineage of SH-SY5Y cells and their established capacity for neurite like process extension in 3D culture [29,30]. Branching began around day 7 (earlier, at day 5-6 in the highest density 15000 group), ultimately consolidating into a stable tripolar morphology by day 12 in the 15000 group including a transient protrusion event on day 11 that retracted and fused back into this configuration, suggesting an architectural preferred, rather than random, endpoint morphology (Figure 1b, Figure S1b). Furthermore, this spatial coincidence of hypoxia and necrosis at branch points suggests that the elongated, protrusion rich architecture of neuroblastoma spheroids creates a local, poorly perfused microenvironment that is distinct from the more uniform hypoxia gradients typically reported in compact spherical tumor spheroid [31].

An early doughnut-shaped cell arrangement was noted transiently on day 1 in 10000 density neuroblastoma group and slightly in the 15000 density glioblastoma group, frequent across densities in optimization trials; this is consistent with an initial ring-consolidation step preceding inward compaction, though we note this interpretation is based on visual observation rather than a validated mechanistic experimentation and should be treated as a working hypothesis instead of an established finding. On other hand, co-culture spheroids grew larger overall than glioblastoma only spheroid (Figure 1c) however retained the spherical morphological characteristics of glioma rather than the elongated architecture of neuroblastoma with all tested cell densities. Although, the 5000 density group alone showing partial deviation from this spherical morphology, suggesting a minimum cell number may be required for stable glioblastoma driven compaction to dominate the co-culture architecture (Figure S1c). Furthermore, necrotic core development in the co-culture was transiently present, but visibly smaller than in neuroblastoma only spheroid, and almost absent in glioblastoma only spheroids system (Figure 1). Nonetheless, co-culture spheroids consistently exceeded diameter ∼ 600 µm across all cell densities, and that spheroids above ∼500 µm are established to develop diffusion limited necrotic cores in line with reported studies [32]. Interestingly, presence of any necrosis in the co-culture despite its glioblastoma like compact architecture indicates that the neuronal component measurably alters the co-culture’s nutrients/oxygen diffusion environment relative to glioblastoma alone, instead of primarily driven by glioblastoma compaction.

As shown in Figure 5, cell tracking data provide insight into architectural differences. For instance, the glioblastoma population within the co-culture suggests increase in relative proportion over time while occupying the least expanded projected area among all groups. This finding suggested that glioblastoma’s growth in 3D environment follow packing of additional cells into a denser structure rather than lateral expansion, thereby making projected area an inverse readout of glioma proliferation in this system. Furthermore, this dense packing tendency appears to be dominant within the co-culture as well, wherein U87-MG displayed its compact growth behavior on the heterotypic culture system, whereas the more loosely self-organizing SH-SY5Y occupies proportionally greater area (Figure 1). Collectively, this finding consistent with glioblastoma’s known capacity to physically and functionally recognize and disseminate into its surrounding brain tissue architecture rather than passively coexist within it [33]. Subsequently, these findings confirm that GBM does not exist as an isolated mass of malignant glial cells but continuously alter the surrounding tissue microenvironment.

Circularity measurements using scale 0-1, wherein 1 indicates highly circular spheroid (Figure 2) further confirmed this architectural distinction quantitatively. Both U87 MG and co-culture spheroids consistently scored between 0.7 and 0.9, whereas SH-SY5Y spheroids largely fell below 0.7 scored and reducing further once branching begun after day 4. For this study, area (instead of volume) and circularity (instead of sphericity) were chosen as the morphological metrics because SH-SY5Y’s elongated and variable depth protrusions (Figure 1b) make extrapolation unreliable. Attempting to infer volume and sphericity from a structure with asymmetric Z-stack protrusions risks introducing systematic morphological misinterpretation rather than providing meaningful biological insight.

**Figure 2.**
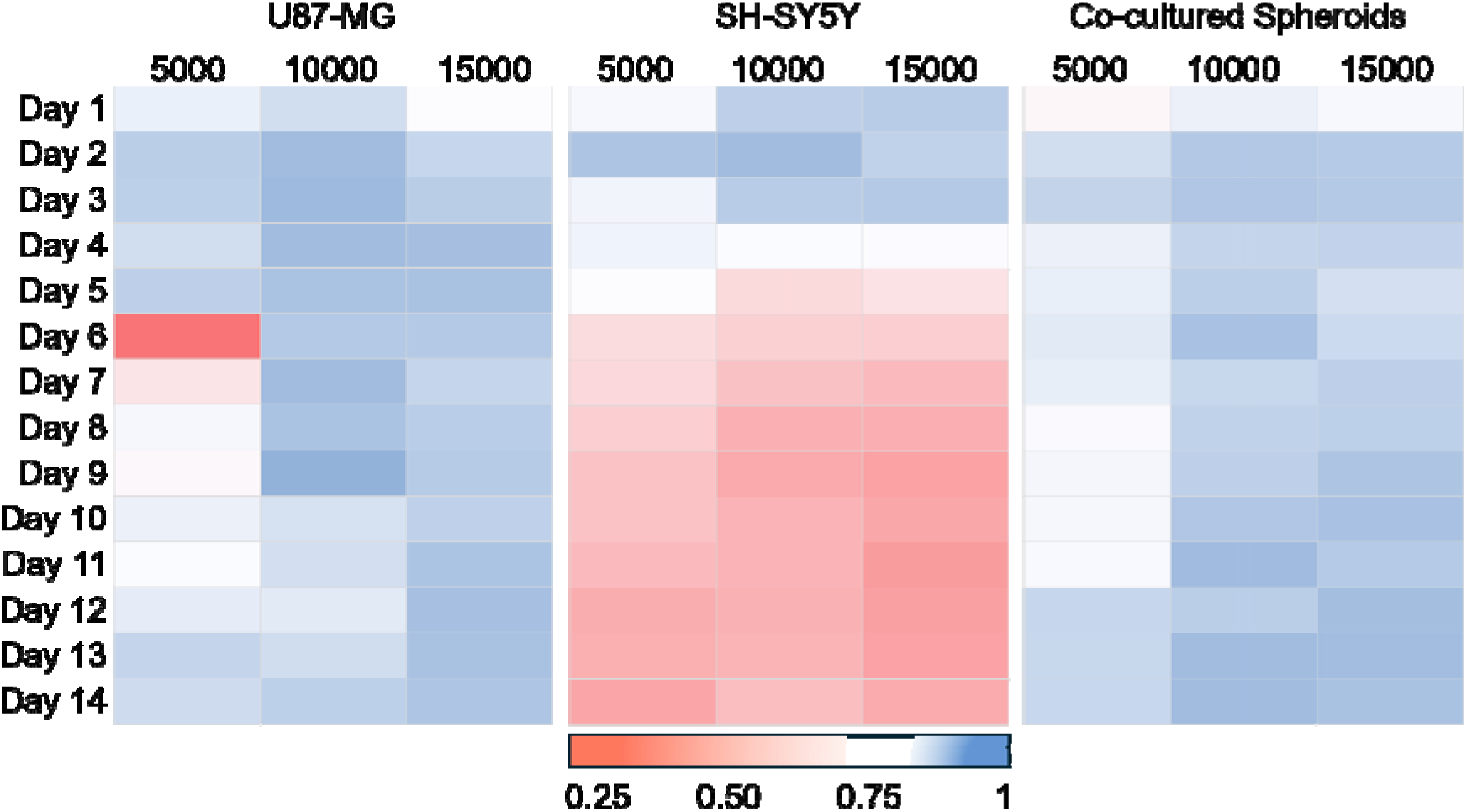
Circularity of Spheroids (n=3) with cell seeding densities of 5000,10000 and 15000 cells: U87-MG (a), SH-SY5Y (b) and co-cultured spheroid (U87-MG)+(SH-SY5Y)) (c).

Furthermore, we performed the growth-rate analysis over the 14 days period (Figure 3) to understand whether the neuronal co-culture alters glioblastoma spheroid development and whether initial cell density influence spheroid architecture and expansion. Notably, monitoring growth over window of 14 days enabled assessment of cell-type specific proliferation, spheroid compaction, and structural architecture, thereby identifying whether neuron-glioma co-culture spheroids adopt a distinct growth phenotype or remain dominated by glioblastoma cells. As shown in figure 3, all the groups showed a common four day spheroid formation window followed by steady growth. The growth rate was clearly cell type specific that is slowest in U87-MG, fastest in SH-SY5Y, and intermediate but skewed toward the slower glioblastoma phenotype in the co-culture system. Interestingly, lower density groups expanded proportionally more across all three spheroid types, probably due to greater per cell nutrient and oxygen availability at lower starting density. We further quantified this expansion over 14 days to validate this pattern. As demonstrated in Figure 4, the densest U87-MG spheroid (density 15000) expanded only ∼10 % compared to ∼40 % at lower densities, while SH-SY5Y expanded far more aggressively (300-600 % across densities) and the co-culture remained consistently intermediate (66-200 %). This density dependent and cell type specific expansion behavior together with co-culture consistently bias toward glioblastoma’s slower denser growth mode. These findings support the conclusion that glioblastoma exerts a dominant architectural influence over the heterotypic spheroid, while neuroblastoma measurably shifts its internal diffusion and necrosis profile.

**Figure 3.**
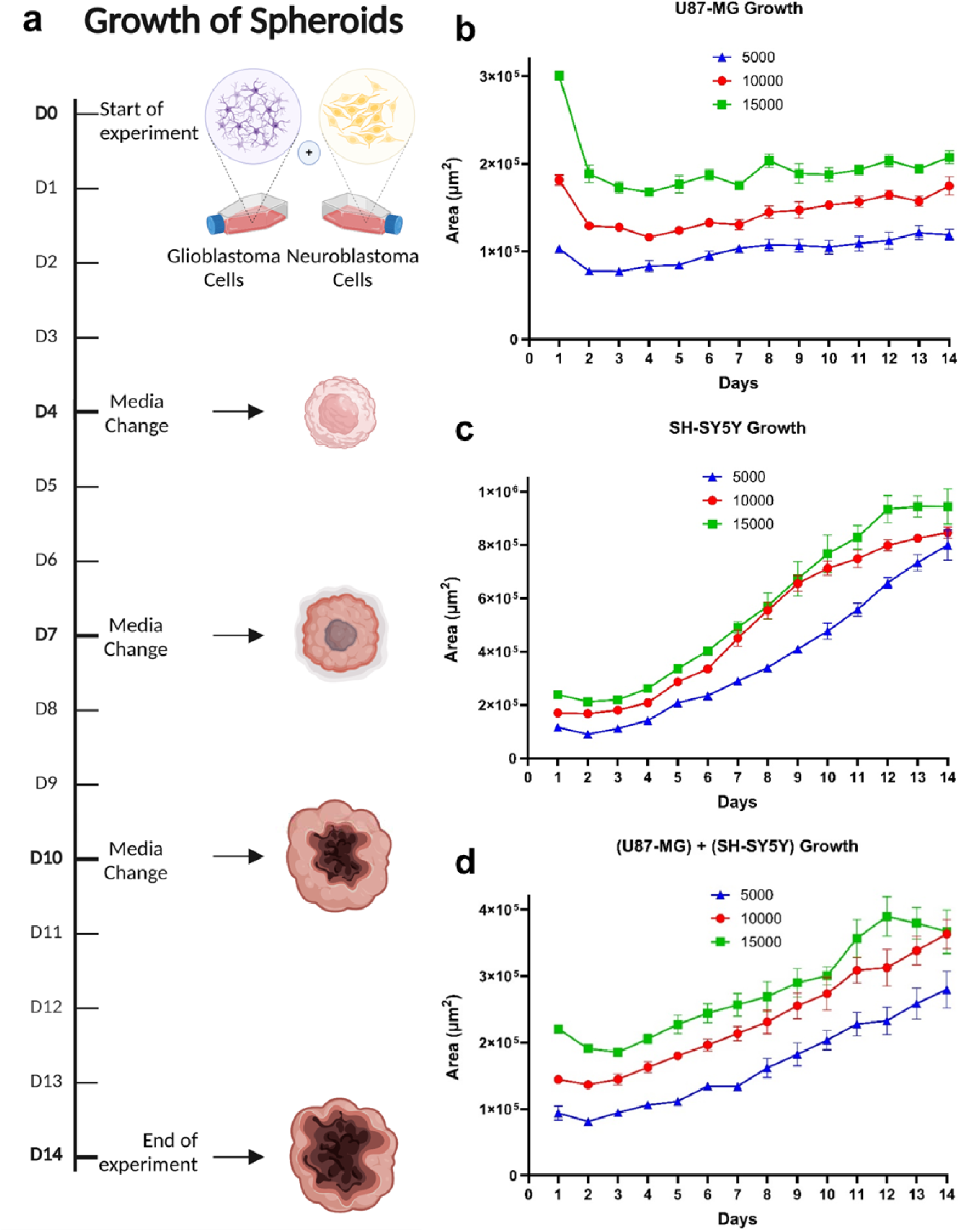
Growth Rate of Spheroids (n=3): Schematic (a) [Created with Biorender.com], U87-MG (b), SH-SY5Y (c), (U87-MG) +(SH-SY5Y) co-cultured spheroids (d)

**Figure 4.**
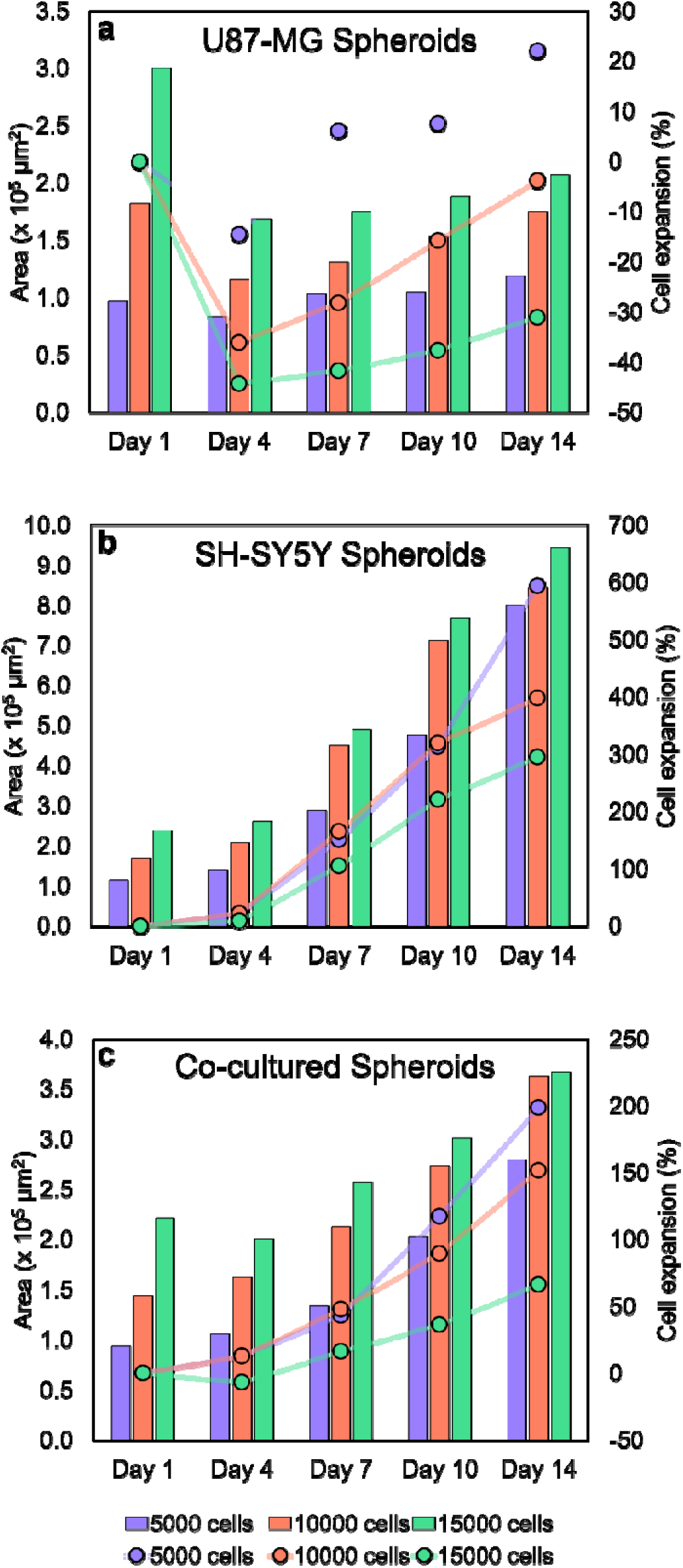
Spheroid Expansion rate: U87-MG (a), SH-SY5Y (b), (U87-MG)+(SH-SY5Y) co-cultured spheroids (c)

**Figure 5.**
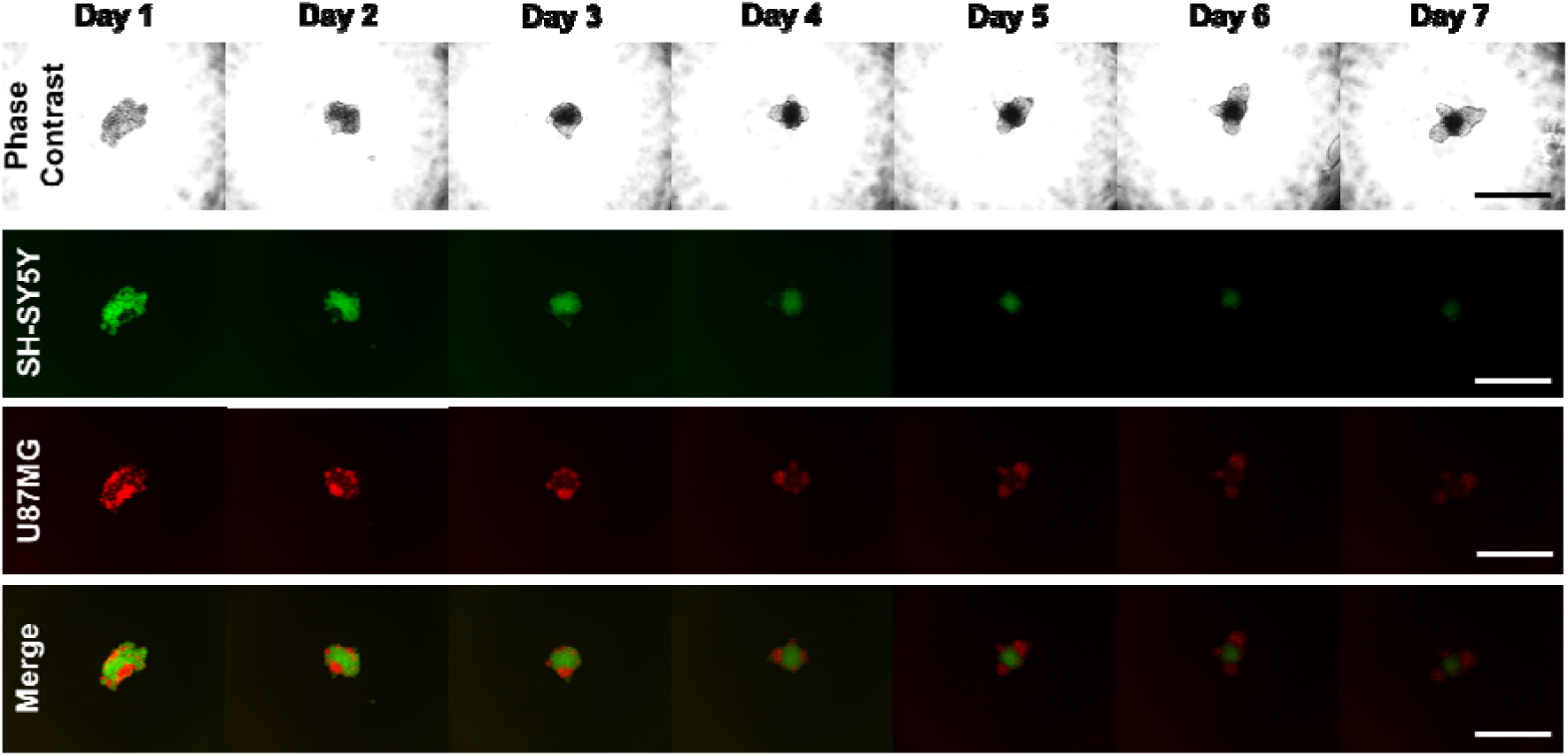
Cell Track Assay performed over 7 days. Scale: 500 μm.

Together, these morphological and growth studies establish that the glioblastoma-neuroblastoma co-culture adopts a predominantly glioblastoma like compact, low necrosis architecture. The same architectural phenotype independently associated with restricted drug penetration and chemoresistance in 3D tumor model consistent with previous studies [34]. The use of a reproducible neuronal surrogate allowed us to investigate whether neuron-glioblastoma interactions alter spheroid architecture and drug response without the technical variability associated with primary neuronal cultures. Based on architecture alone, this would predict that the co-culture should resist temozolomide (TMZ) similar to glioblastoma only spheroid. Whether the presence of neuronal lineage component however alter this expected drug response as our central hypothesis is addressed directly in the following section.

### 3.2. Spatial Organization

Having established in Section 3.1 that the co-culture adopts a predominantly glioblastoma like compact architecture, we next investigate whether this reveals true intermixing of the two cell populations or, instead, a spatially organized arrangement within the spheroid. These two possibilities carry different implications for how neuronal cues could possibly influence glioblastoma growth and behavior. As shown in Figure 5 fluorescence cell-tracking of glioblastoma (red) and neuroblastoma (green) over 7 days determined this directly wherein the two populations did not form a homogenous mixture but instead segregated into distinct spatial compartments. At seeding, glioblastoma cells initially dispersed randomly followed by progressively redistributed toward the spheroid periphery and consolidated into outer-rim clusters, whereas neuroblastoma cells became confined to the spheroid core. The gradual decrease in fluorescence intensity across the 7 day window is consistent with dye dilution through successive cell division rather than cell loss and the comparatively greater retained fluorescence in the glioblastoma population over time consistent with its established higher proliferative or doubling rate relative to SH-SY5Y (Section 3.1). This systematic core versus rim discrimination is functionally significant. Compartmentalized spatial organization of this kind has been reported in other heterotypic spheroid systems as a consequence of differential cell-cell adhesion strength and proliferative pressure. For instance, the more cohesive and faster growing population preferentially occupies the outer, better perfused rim while the less cohesive population is displaced inward [35,36]. This agreement places the neuronal lineage surrogate directly within the spheroid’s most oxygen and nutrient restricted zone, which is the same core region established in Section 3.1 as the site of hypoxia onset and eventual necrosis in neuroblastoma only spheroids. This is a crucial structural anchor for interpreting the drug response such as temozolomide attributed to the neuronal component in cells positioned at the spheroid’s pharmacologically hard to penetrate and most microenvironmentally stressed region, not cells freely intermixed with glioblastoma at the drug accessible rim. Thus, this spatial configuration will be directly relevant to interpreting both the hypoxia and proliferation/viability outcomes and the temozolomide sensitivity results that follow.

### 3.3. Viability and necrotic core development

Building on this spatial architecture, we next assess the viability qualitatively using FDA/PI dual staining on day 7, day 10 and day 14 (Figure 6) to determine whether the core-confined neuronal population and rim occupied glioblastoma population show correspondingly distinct survival behaviors. FDA is hydrolyzed by intracellular esterases in live and membrane intact cells to fluorescein (green), whereas PI enter only membrane compromised cells and intercalates with DNA to emit red fluorescence, tagging the dead cell population. Across all three time intervals, glioblastoma viability remained consistently high, while neuroblastoma viability progressively declined. This characteristic corroborates glioblastoma’s known capacity to tolerate hypoxia, nutrient restricted conditions and sustain proliferative activity under microenvironmental stress that is otherwise cytotoxic to less resilient cell type [37]. Within the co-culture system, PI staining was concentrated in the spheroid core the same region identified in Section 3.2 as the neuroblastoma occupied compartment. These findings support the interpretation that the observed core necrosis reflects neuroblastoma cell death rather than glioblastoma death within the heterotypic structure.

**Figure 6.**
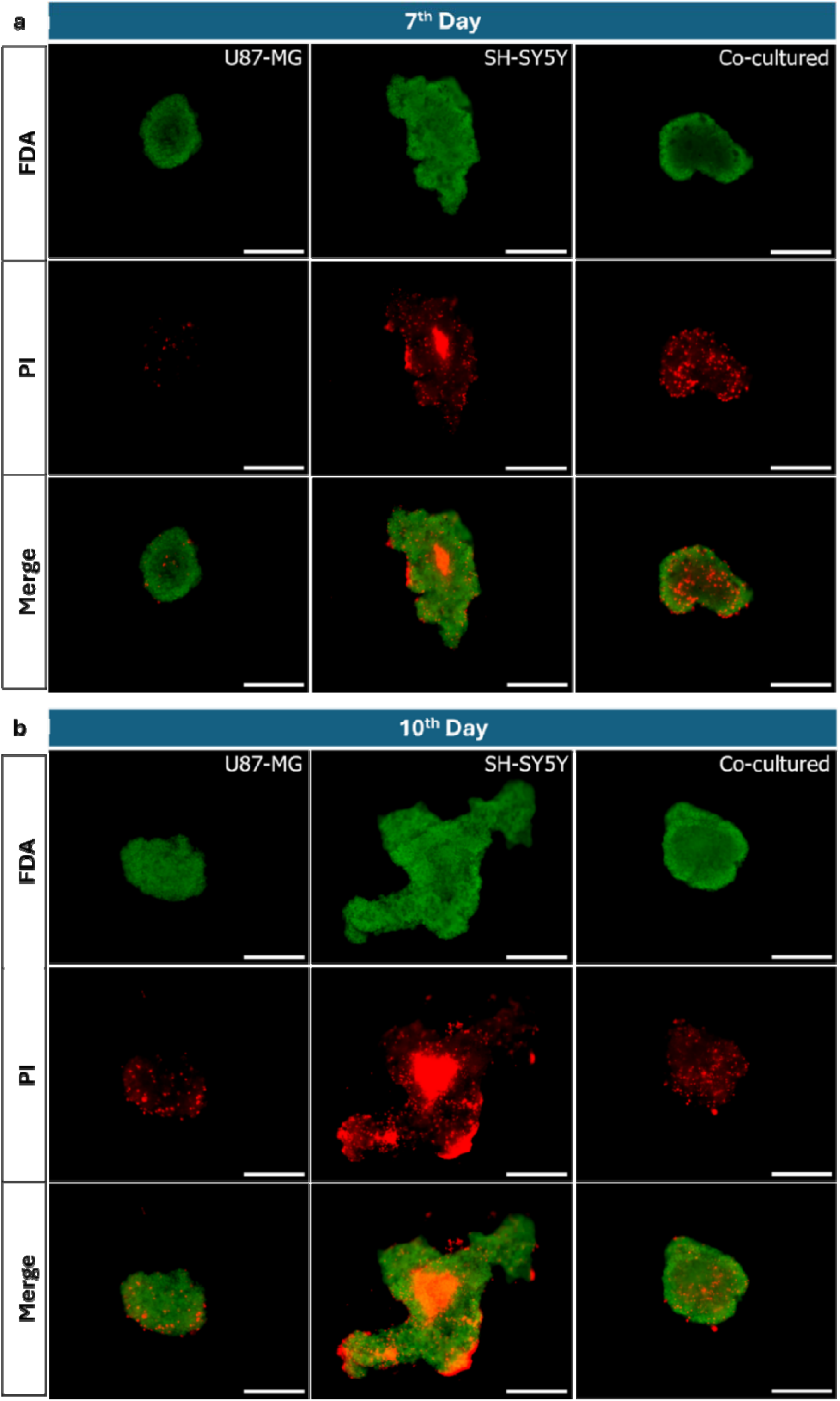
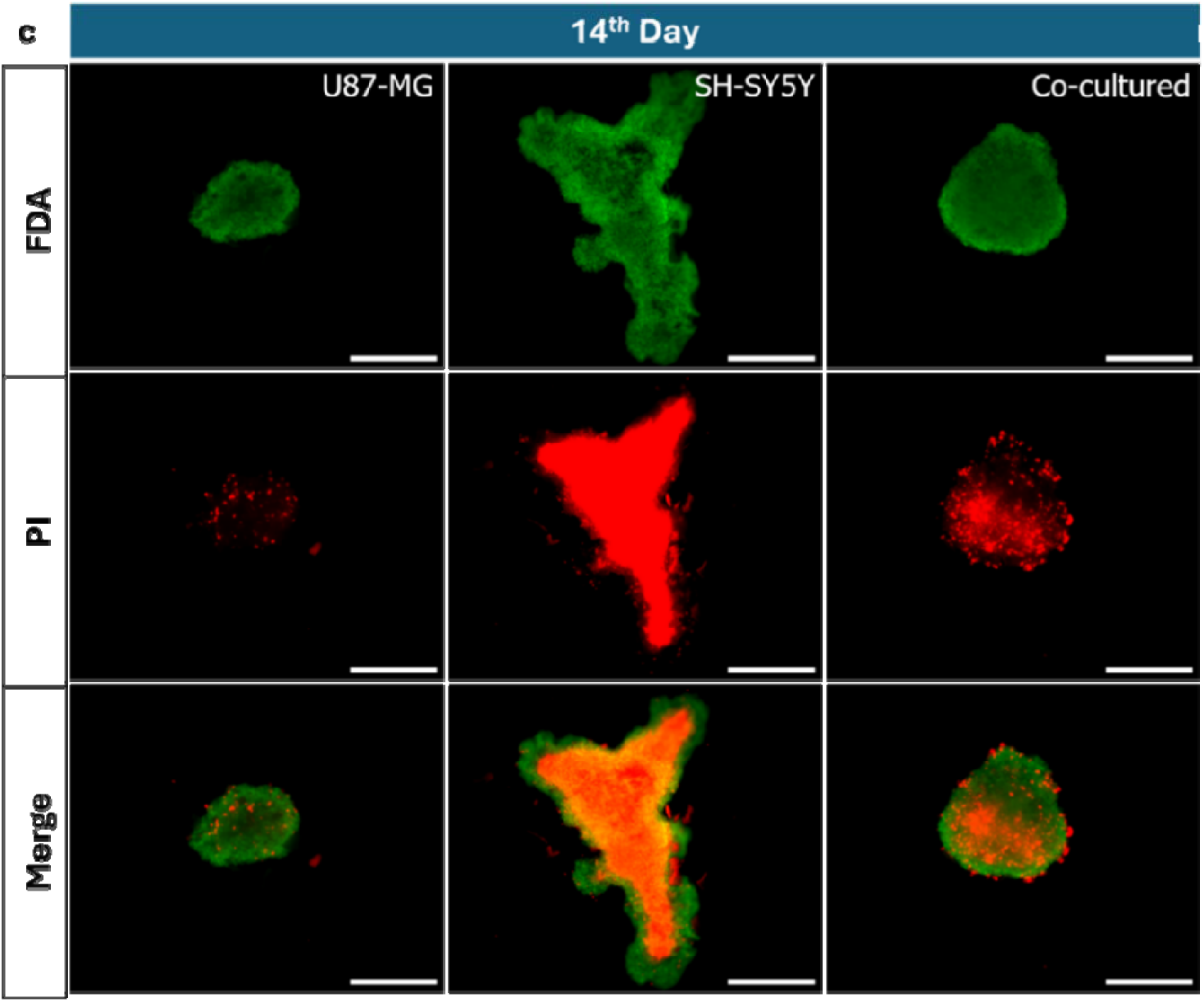
Live/Dead results - Day 7 (a) Day 10 (b) Day 14 (c) Scale 500 µm.

Mechanistically, we attribute this differential viability primarily to packing geometry rather than an intrinsic difference in hypoxia tolerance alone. As established in Section 3.1, neuroblastoma’s loosely packed and large area architecture is inherently less favorable for nutrient and oxygen penetration than glioblastoma’s dense compaction, amplifying whatever intrinsic hypoxia sensitivity SH-SY5Y cells may have. Collectively, this finding establishes a coherent structural feature i.e. glioblastoma’s rim localized, compact, well tolerated growth physically and functionally marginalizes the neuronal lineage population into a core zone that is simultaneously hypoxic, poorly perfused, and progressively necrotic. Interestingly, this raises a direct mechanistic question whether a neuronal population under this microenvironmental stress can still exert a measurable influence on glioblastoma’s drug response, or whether its confinement and gradual death effectively silence its contribution to the co-culture’s overall chemosensitivity.

### 3.4. Hypoxia precedes necrosis

Distinct rim versus core spatial compartment within the co-culture of glioblastoma and neuroblastoma respectively further correlates with different cellular viability outcomes in which glioblastoma remain highly viable, and neuroblastoma progressively necrotic. Therefore, to understand whether hypoxia is the mechanistic link connecting spatial organisation to this differential fate, dual staining with PI and hypoxia responsive probe was performed on day 7, distinguishing regions of oxygen deprivation (< 5 % O_2_) from dead cells within the same spheroid (Figure 7).

**Figure 7.**
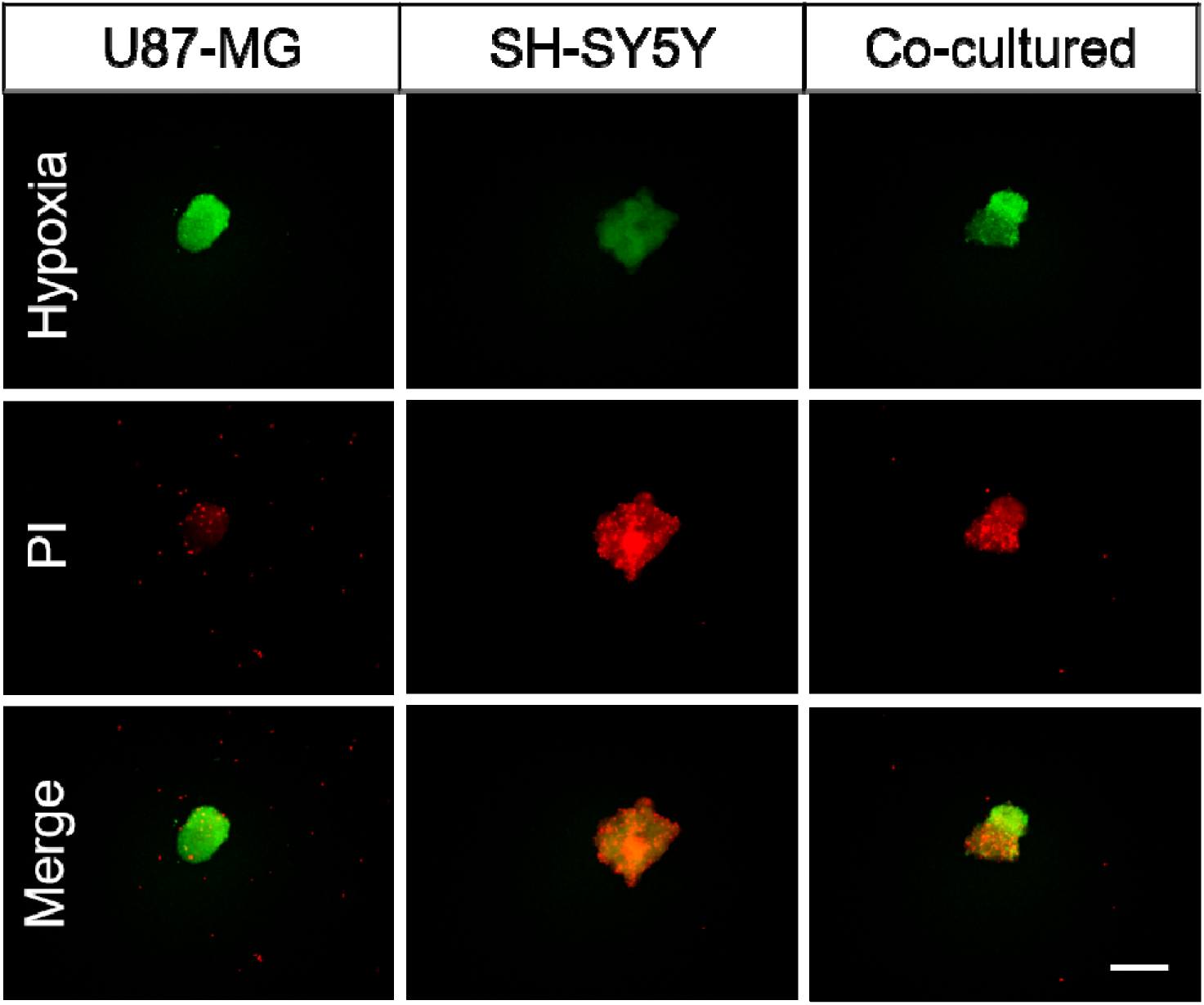
Hypoxia Imaging on Day 7. Scale 200 µm.

Extensive hypoxia was found throughout the spheroid in all three groups, indicating that oxygen limitation itself is not what distinguishes glioblastoma from neuroblastoma spheroids, as both experience substantial hypoxic stress on day 7. Furthermore, localization and extent of PI positive cells relative to the hypoxic zone differentiates the groups. In glioblastoma only spheroids, the merged image showed minimal PI intensity despite extensive hypoxia, suggesting that glioblastoma cells largely survive under oxygen deprived condition. Thus the hypoxia and cell death relation is consistent with glioblastoma’s well-established capacity to exploit hypoxic signaling for tumor progression and recurrence [38]. Similarly, as evidenced by previous studies hypoxia inducible factor HIF-1α is a key driver of glioma stemness, proliferation, and adaptation within the tumor microenvironment [39] and the minimal cell death despite confined oxygen deprivation here is consistent with that adaptive association to hypoxia. Though the mechanism of action of the hypoxia marker used is not disclosed by the manufacturer, considering the strong relation between hypoxia and HIF-1α, the above said is strongly plausible.

Conversely, neuroblastoma only spheroids showed the opposite behavior i.e. extensive cell death forming a necrotic core, spatially consistent with the necrotic region observed in the day 7 live/dead cell assay (Section 3.3., Figure 6a). This finding indicates that unlike glioblastoma, neuroblastoma’s hypoxic exposure influence the cell death rather than adaptive survival consistent with the packing density driven diffusion constraint proposed in Section 3.3. Neuronal lineage cells generally lack the hypoxia adaptive programs such as HIF-1α drive stemness or proliferative signaling that protect glioma cells under equivalent oxygen restriction.

In the co-culture spheroid, the merged hypoxia and PI image revealed two spatially distinct zones such as one that predominantly hypoxic but PI negative and other predominantly necrotic. Based on the spatial distribution established by cell tracking study, together with each monoculture’s independent hypoxia responsive phenotype established above, the most promising interpretation could be that the hypoxia but viable zone corresponds to the glioblastoma occupied region, whereas the necrotic zone corresponds to the neuroblastoma occupied core. Collectively, this establishes a reasoned structural preclinical model of GBM co-culture in which glioblastoma occupies the rim in a hypoxic but non-necrotic, adaptively thriving state, while neuroblastoma is confined to a hypoxic and necrotic core. This finding is significant for the subsequent drug-response studies. It indicates that the neuronal lineage is not only present but also undergo hypoxia-induced cell death by the time of temozolomide treatment is assessed. Furthermore, this raises the question of whether the remaining neuronal population still influence glioblastoma chemosensitivity, or whether the observed differences in drug response are predominantly due to neuronal cell death than specific neuron-glioblastoma interactions.

### 3.5. Temozolomide response

As the co-culture adopts a glioblastoma driven architecture (compact, rim localized and highly hypoxia tolerant) while confining a progressively dying neuroblastoma population to its necrotic and hypoxic core. Based on this architecture and consistent with the established link between drug resistance and compact, poorly perfused 3D structure indicated that the co-culture would be predicted to show temozolomide resistance similar to glioblastoma only spheroid system. Testing this hypothesis will help us to predict alteration in temozolomide sensitivity in presence of neuronal lineage cells, and independent of what its architecture alone. Chemosensitivity was initially screened using the MTT assay across a range of concentration, but results as shown in Figure S3 were inconsistent. This is an expected limitation of a metabolic penetration dependent colorimetric readout in a 3D structure where dye access is itself confounded by the same diffusion barriers under investigation. Moreover, luminescent ATP-based endpoint assays overcome this limitation by lysing the whole spheroid and reporting total ATP in direct proportion to viable cell number, rather than depending on differential dye penetration. This choice is further supported by proof that tetrazolium based (e.g. MTT assay) absorbance correlates reliably with viable cell number only across a limited spheroid range. While ATP-based luminescence endpoint correlates linearly across diameters from 100-1000 µm [40] spanning the full size range observed in this study. Consistent with this, CCK-8 a related colorimetric assay applied to 3D co-culture of U87-MG and SH-SY5Y, however it has also failed to yield a valid cytotoxic readout [41]. Therefore, Cell-Titer Glo 3D an ATP-based assay was selected for temozolomide effectiveness screening. Across all three spheroid groups temozolomide was given on day 7 as a single dose (10, 100 and 1000 µM) and the viability was assessed after 24 h to enable direct and matched comparison (Figure 8).

**Figure 8.**
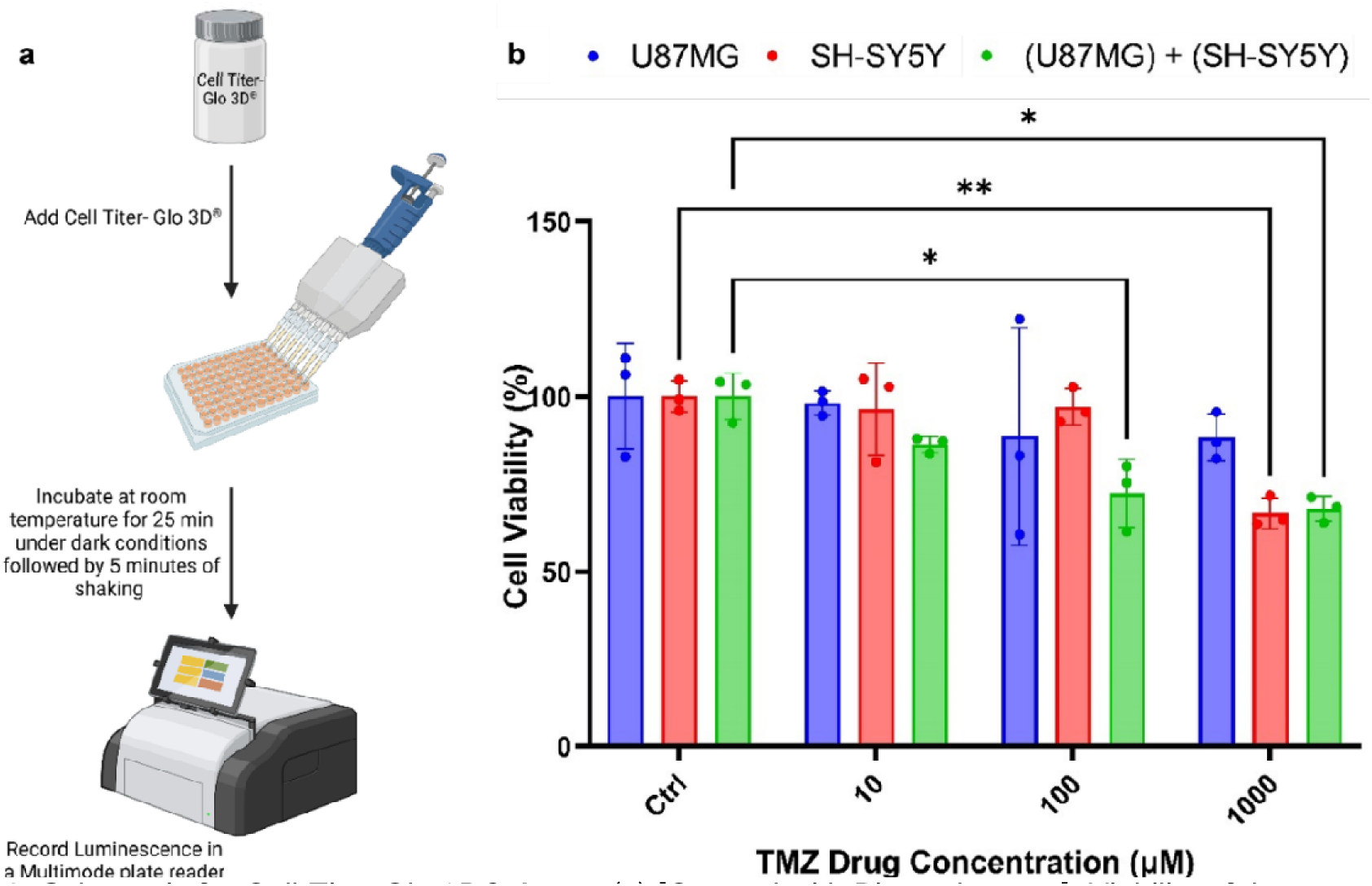
Schematic for Cell-Titer Glo 3D® Assay (a) [Created with Biorender.com], Viability of the spheroids on Day 8 (after 24h DT on Day 7) (b). n=3. *p<0.05, ** p<0.01.

We found that at 1000 µM, glioblastoma only spheroids retained ∼88 % viability showing resistant across the entire tested range. While neuroblastoma only and co-culture spheroids fell to ∼ 67 % and ∼ 68 % respectively. Neuroblastoma only spheroid became sensitive above 100 µM dose. The co-culture, by contrast responded at every concentration tested, with viability declining progressively from 86 % to 72 % to 68 % across 10, 100, and 1000 µM doses. Interestingly, at 10 µM neither monoculture responded at all (both showed viability ∼ 97 %), though the co-culture already showed a clear reduction, demonstrating that co-culturing confers a low-dose sensitivity absent from either parent population alone, and consistent with the study’s central hypothesis that the neuronal lineage component does not passively co-exist with glioblastoma but measurably alters its drug sensitivity, in a direction opposite to what compact and hypoxia-tolerant architecture alone would predict.

Morphological changes accompanying temozolomide treatment (Figure S4 & Figure S5) demonstrated that glioblastoma only spheroid area decreased at 10 and 100 µM and further at 1000 µM doses, with disaggregated small cell clusters appearing in phase-contrast images at the highest concentration. A disaggregation patterns also found across the other two spheroid groups at 1000 µM. Neuroblastoma only spheroids continued to expand in area under control, 10 µM and 100 µM conditions, but showed no further expansion at 1000 µM. This findings are in agreement with temozolomide suppressing SH-SY5Y proliferation rather than inducing shrinkage based cell death at this timepoint. Although, co-culture spheroids showed no significant area change, however did show a clear concentration dependent trend with reduced area expansion at 10 µM relative to control, no change at 100 µM, and a net area reduced at 1000 µM post 24 h treatment. Nonetheless, a graded response was observed across all three tested concentrations which would not be expected if the effect were driven solely by a static, already dead core population than an active, dose responsive process. Dead cell within the co-culture spheroids likely remain structurally retained possibly due to the mechanical integrity conferred by the endogenous, self-assembled extracellular matrix (ECM) that forms in scaffold-free spheroid culture [42].

As shown in Figure 9 live/dead cell assay performed on day 8 (24 h post temozolomide treatment) corroborated the ATP-assay findings closely. Glioblastoma only spheroids remained predominantly FDA-positive across all concentrations, with minimal PI staining. At 1000 µM treatment, loosely bound peripheral cells were displaced consistent with the size reduction noted above. However, these displaced cells remained viable, matching ∼ 88 % viability retained even at the highest TMZ concentration in the ATP assay. Neuroblastoma only spheroid showed elevated baseline cell death even in the untreated control, with FDA-positive signal restricted primarily to the proliferative zone. The 10 and 100 µM conditions demonstrated negligible changes relative to control, once again mirroring the ATP-based assay pattern, while 1000 µM showed near complete PI dominance. The fluorescence readout therefore indicated a more complete loss of viability at 1000 uM than the ∼67% retained in the ATP assay, reflecting the different endpoints measured by the two methods. In contrast, in co-culture both the qualitative and quantitative assays were fully consistent. A clear dose dependent response to chemotherapy was observed from 10 µM onwards, which was not seen in either monoculture. At 1000 µM, the displaced cells in the co-culture were completely PI positive indicating cell death, while the displaced glioblastoma cells in the monoculture remained largely viable at the same dose. This finding suggest that glioblastoma responds differently in the co-culture than in monoculture. The co-culture appears to improve glioblastoma sensitivity to temozolomide induced cell death. Further studies are needed to confirm whether this effect is caused from neuron-glioblastoma interactions rather than neuroblastoma cell death alone.

**Figure 9.**
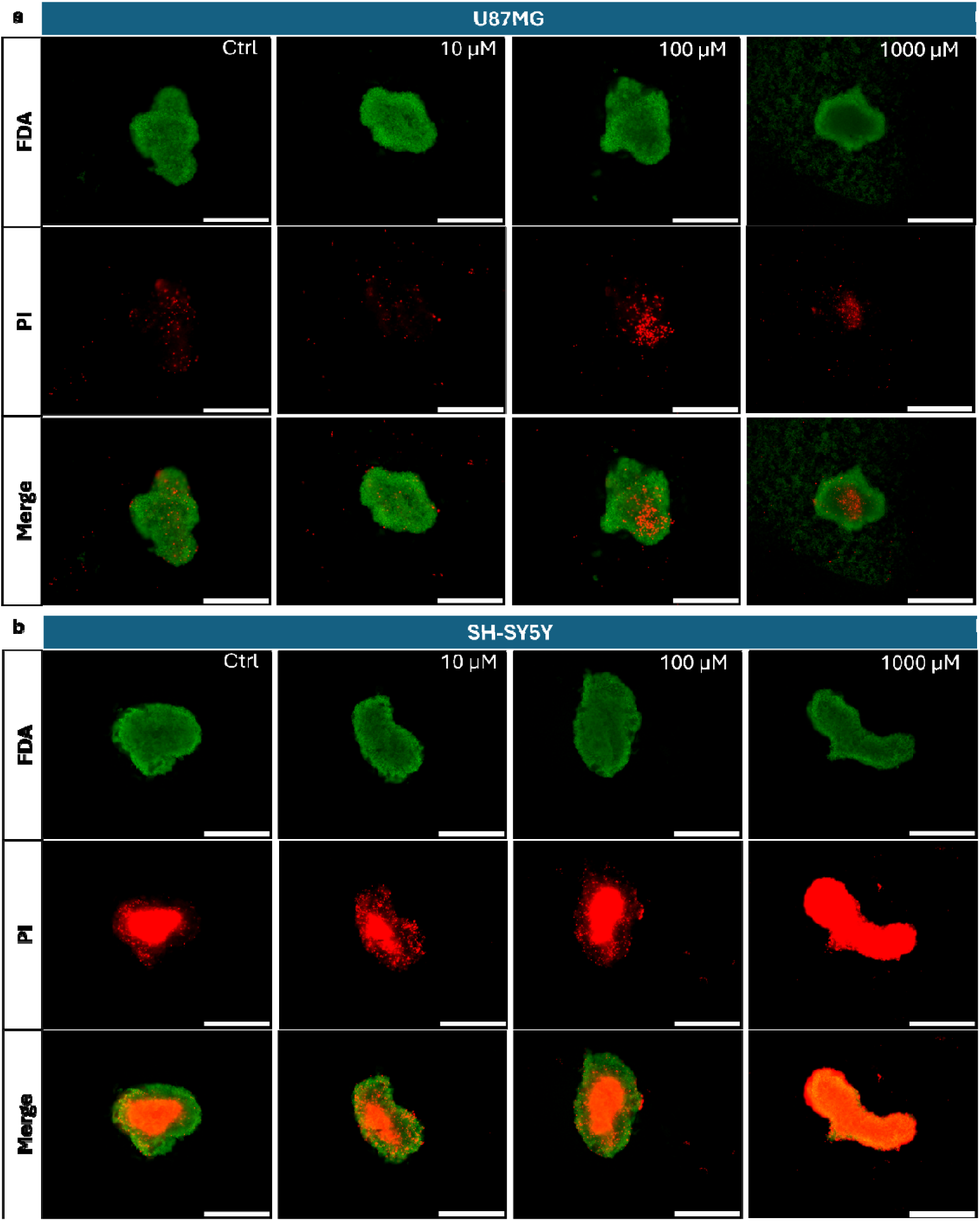
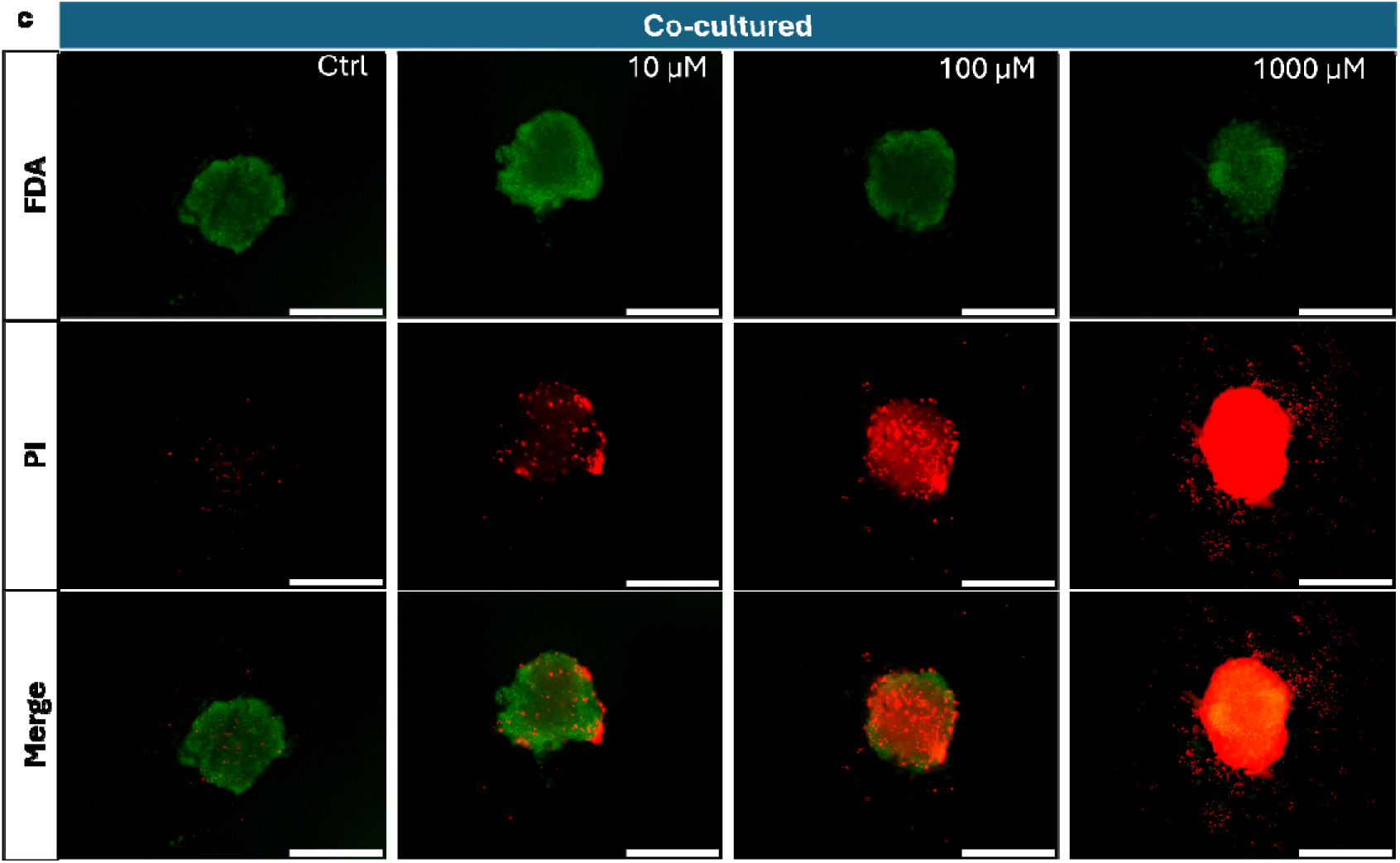
Live/Dead Assay on the spheroids on Day 8 (after 24h DT on Day 7)

The resistance observed in glioblastoma only spheroids likely reflect an intrinsic property of the 3D structure rather than the cells alone. Previous studies have reported the IC_50_ of temozolomide at ∼124 µM after 24 h on the 2D glioblastoma (U87-MG) monoculture, albeit with considerable inter-study variability [43,44]. Wherein in 3D culture, even 1000 µM was not sufficient to reduce viability by 50 %. This is consistent with the previous studies showing that drugs are generally less effective in 3D spheroid models than in 2D monolayer cultures [45,46], and that the response of glioblastoma to temozolomide depends on the experimental model and culture choice [47,48]. Key structural factors may contribute to this effect. The compact spheroid architecture can limit drug penetration [49], while the hypoxic core reduces the effectiveness of temozolomide, whose mechanism depend on actively proliferating cells to induce DNA damage [50,51]. In addition, the self-produced extracellular matrix in scaffold free spheroids may further restrict drug response, with collagen rich matrix reported to reduce drug efficacy in 3D spheroid models [42,52,53]. Further mechanistic studies are needed to determine the relative contribution of each factor.

Moreover, our findings also showed that spheroid morphology alone does not predict drug response. Although, glioblastoma only spheroids showed high circularity, they were the most resistant to temozolomide among all three groups. This contrasts with the study by Senrung et al; which suggested that higher spheroid circularity is associated with greater drug sensitivity [21]. These results indicate that spheroid circularity alone is not a reliable predictor of chemosensitivity and support the use of functional drug response assays, rather than morphology alone for evaluating 3D glioblastoma models.

The most significant finding of this study is that the co-culture spheroids responded to temozolomide even at 10 µM, whereas neither the glioblastoma only and neuroblastoma monoculture spheroids showed a response at this concentration. These results were consistently observed in both the live/dead and Cell Titer Glo-3D assays. These findings support the use of a neuron-containing 3D glioblastoma model than a glioblastoma only model, as glioblastoma is known to interact closely with neurons during tumor progression [8,11]. Our results suggest that the presence of neuronal cells influence the drug response of glioblastoma, underscoring the importance of incorporating relevant cells from the tumor microenvironment into preclinical drug screening models.

This study provides a translational proof-of-concept for a neuron containing glioblastoma spheroid model but does not address the underlying mechanisms. Future studies should investigate the molecular basis of the altered temozolomide response, including whether it is driven by direct neuron-glioblastoma interactions, metabolic effects, or the contribution of neuronal cell death. In addition, the spatial cell tracking and hypoxia observations should be validated using quantitative biomarker based analyses. Such studies will further strengthen the utility of this co-culture model for preclinical drug screening.

## 4. Conclusion

The present study establishes a heterotypic 3D co-culture model combining glioblastoma (U87-MG) and a neuronal lineage population (SH-SY5Y) developed to understand previously unattended question whether the presence of neuronal cells directly alter glioblastoma’s response to temozolomide in a scalable 3D spheroid system independent of tumor vasculature, immune infiltrate, or the extended maturation timelines that restrict organoid based ex-vivo platforms. Spheroids were generated using the liquid overlay method using agarose over 14 days and characterized for growth, morphology, spatial organization, and hypoxic profile.

Glioblastoma spheroids were found to be consistently dense, highly circular, and strongly temozolomide resistant (88 % viability at 1000 µM). Whereas neuroblastoma spheroids were larger, less dense and circular, and chemo sensitive (67 % viability at 1000 µM) with a size-dependent necrotic core. Fluorescence cell tracking and dual hypoxia/viability staining showed that glioblastoma and neuroblastoma reside in distinct spatial compartments within the co-culture i.e. glioblastoma at the rim remaining largely viable despite hypoxia and neuroblastoma confined to a hypoxic, progressively necrotic core. Notably, the co-culture adopted the glioblastoma’s compact and circular architecture. However, this glioblastoma like architecture which based on established links between compact 3D structure and restricted drug penetration, would predict temozolomide resistance alike to glioblastoma alone. The co-culture instead responded to temozolomide across the entire concentration range tested, including at 10 µM (86 % viability), a concentration to which neither monoculture responded at all, ultimately reaching 68 % viability at 1000 µM dose. Our results suggest that the presence of neuronal lineage cells enhances glioblastoma sensitivity to temozolomide, independent of spheroid architecture, underscoring a response that would not be captured by glioblastoma only models. Collectively, these findings highlight the significance of including neuronal cells in preclinical glioblastoma models, as the neuronal microenvironment may influence therapeutic response. This study provides a translational preliminary proof-of-concept for a neuron occupied glioblastoma spheroid model. Future studies should investigate the molecular mechanisms underlying the enhanced temozolomide response and evaluate whether it is directly driven by neuron-glioblastoma interactions or other factors. Nonetheless, to validate the spatial organization and hypoxia findings biomarkers based investigation are needed. Additionally, comparison with patient-derived organoids models will further help to understand the utility of this simplified culture system for high throughput drug screening studies.

## Data statement

All data presented in this manuscript is available from the corresponding author on request.

## Author Contributions

Conceptualization – NT, SM. Data Curation – SM. Formal Analysis – SM. Funding Acquisition – NT. Methodology – SM. Project Administration – SM, NT. Resources – NT. Software – SM, AM. Supervision – NT. Validation – SM. Visualization – SM, AM. Writing/Original Draft Preparation – SM, PS. Writing/Review & Editing – SM, PS, NT.

## Supporting information

GBM-NBM TMZ Supplementary File

## Acknowledgements

Authors acknowledge funding under the Science Foundation Ireland and Irish Research Council (SFI-IRC) pathway program (21/PATH-S/9634).

