## Supplementary material for "Glioblastoma–Neuroblastoma co-cultured multicellular spheroid model for evaluating Temozolomide response": GBM-NBM TMZ Supplementary File

a) **U87-MG**

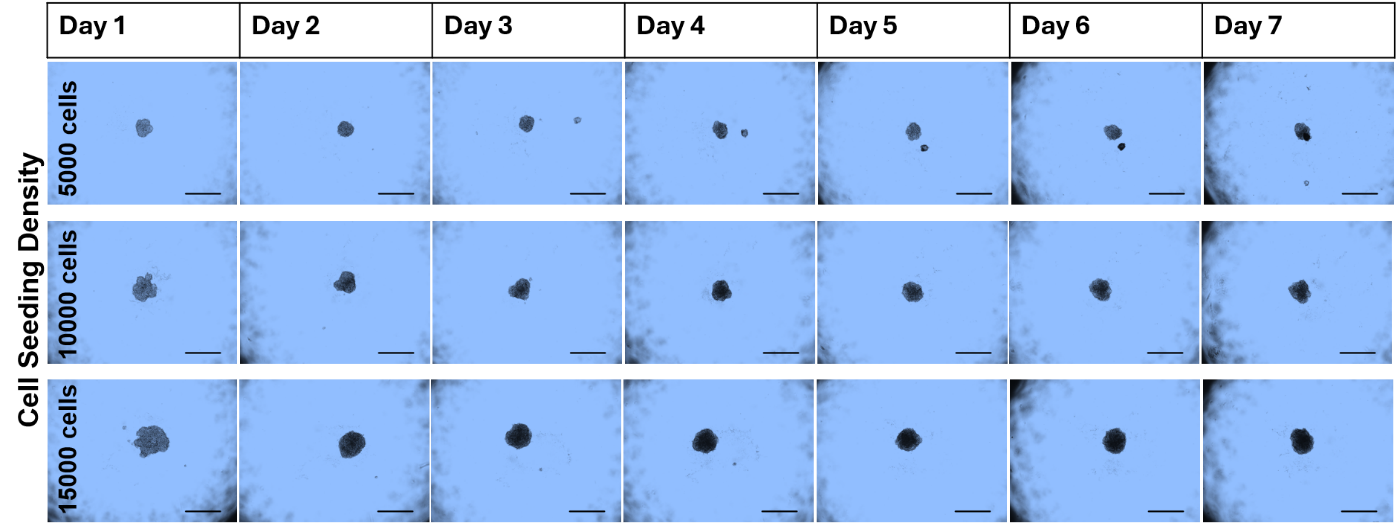

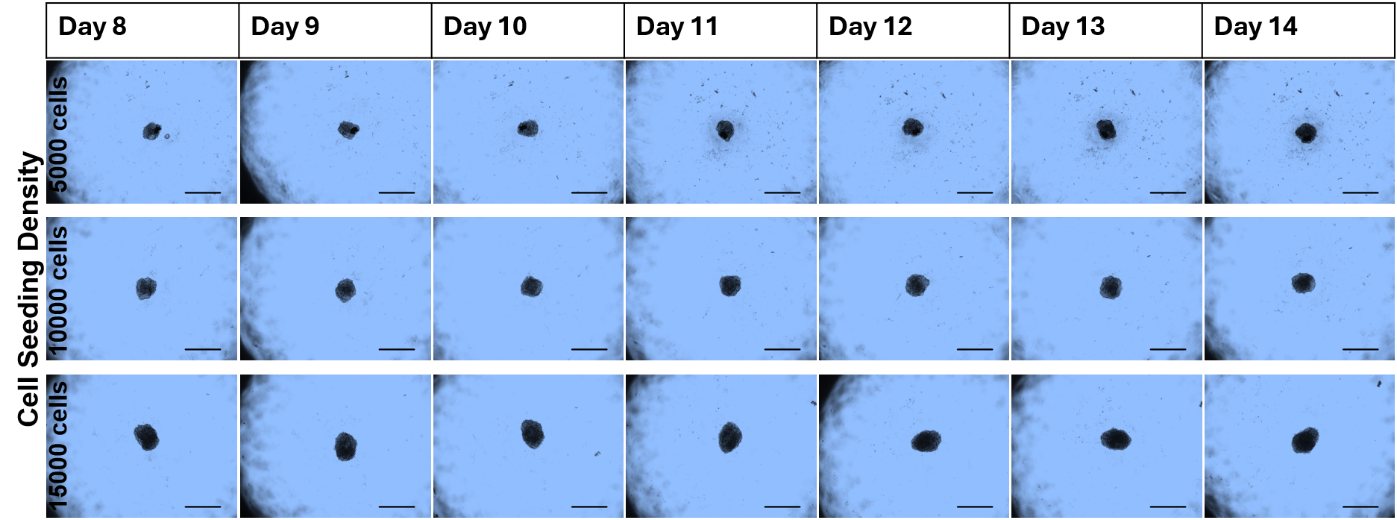

b) **SH-SY5Y**

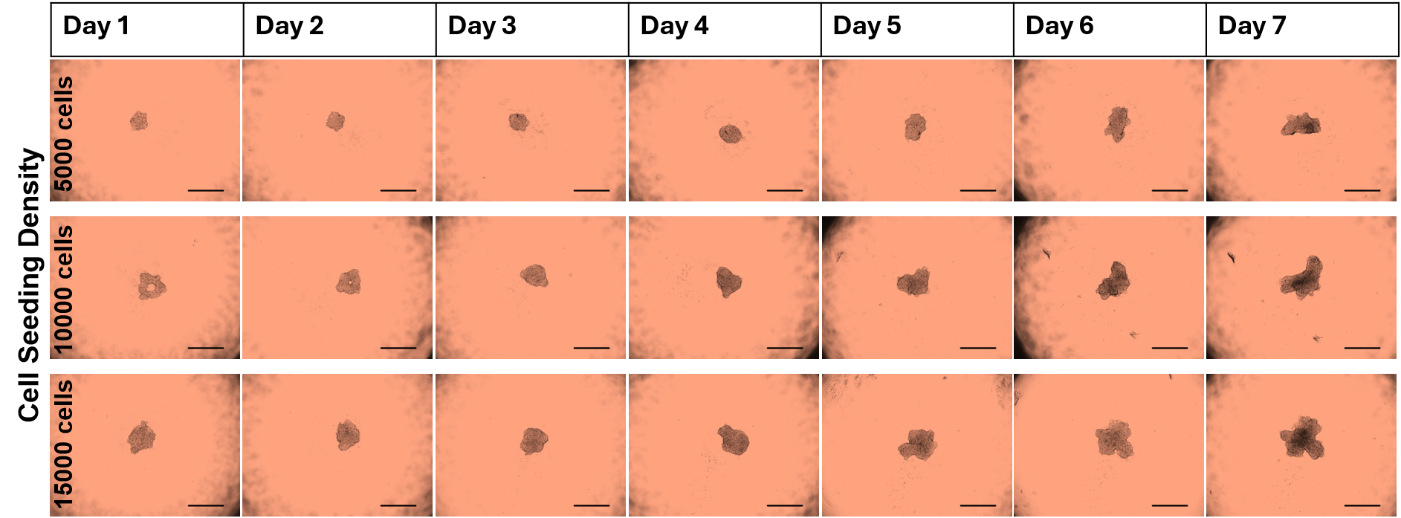

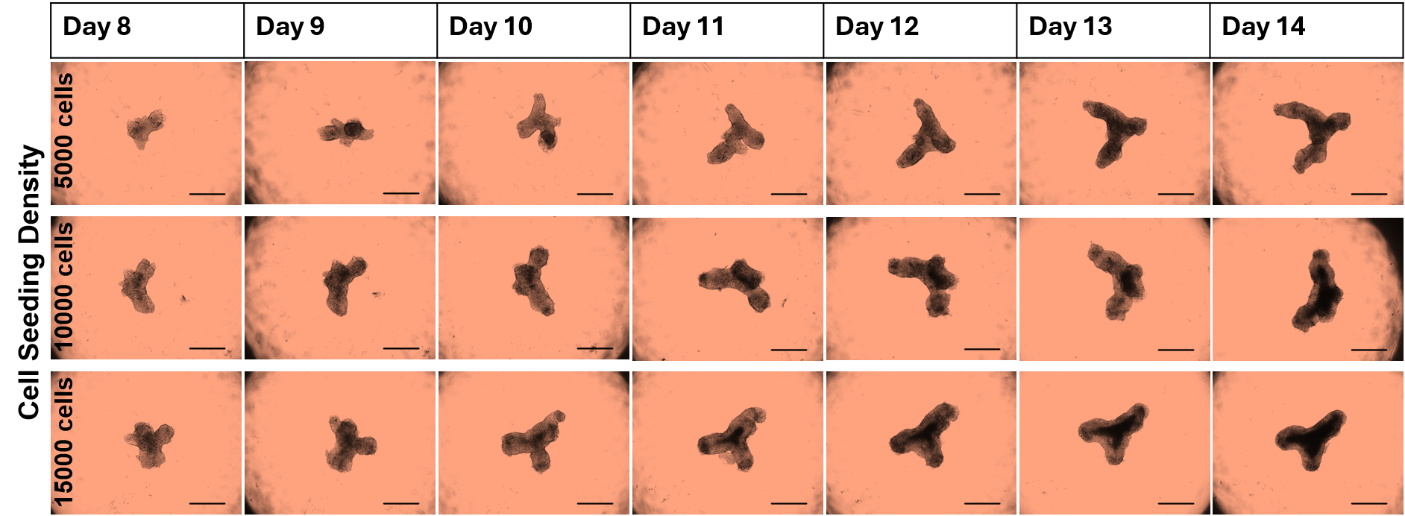

c) **Co-cultured Spheroids**

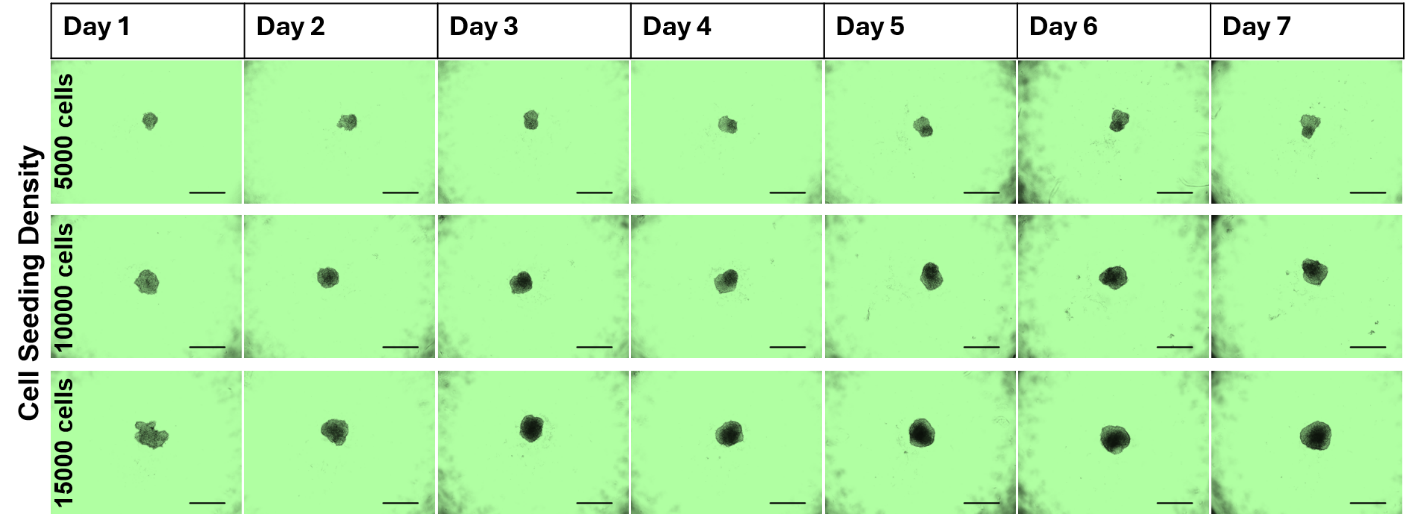

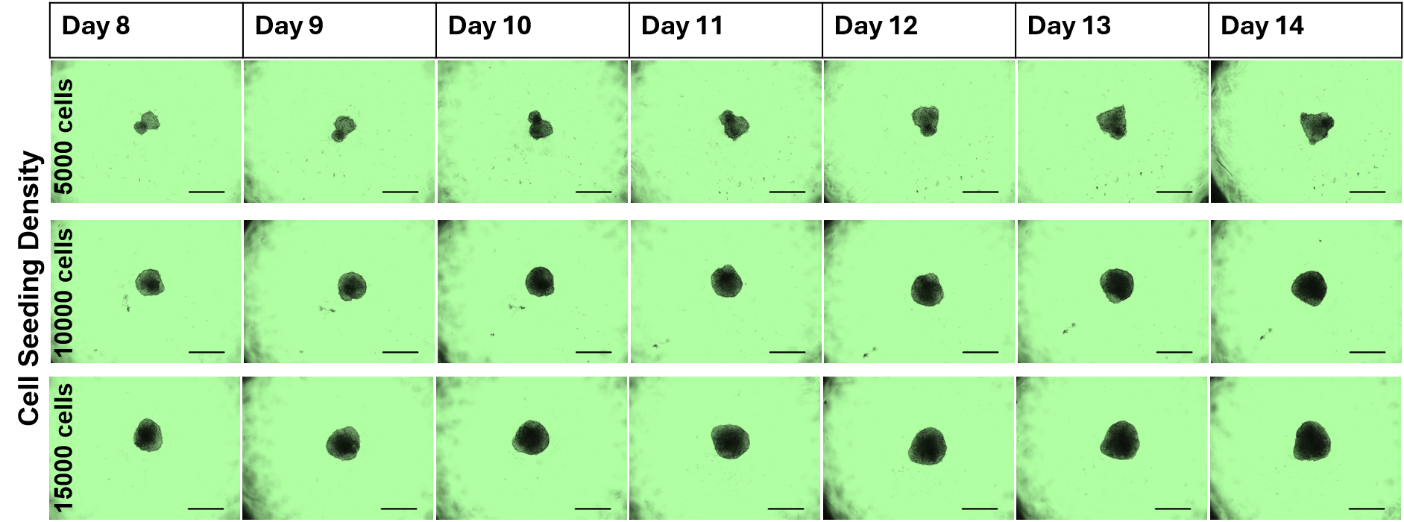

**Fig S1:** All 14 days phase-contrast images of spheroids acquired using 4X objective lens in Keyence BZ-X810 microscope. a) U87-MG Glioblastoma Spheroids b) SH-SY5Y Neuroblastoma spheroids c) Co-cultured Spheroids.

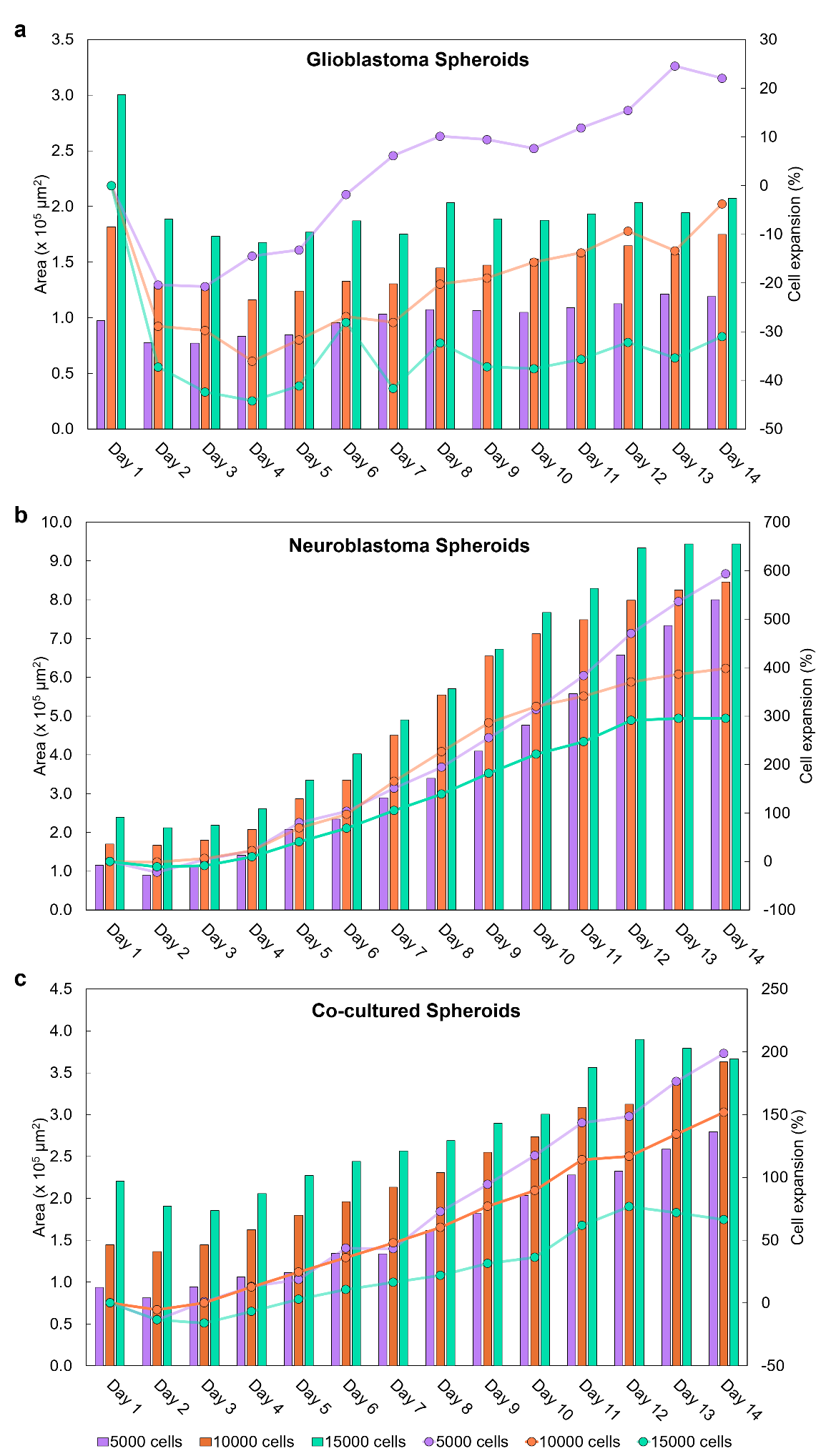

**Fig S2:** Area and Percentage of area expansion in spheroids grown with 5000 to 15000 cell seeding densities over a period of 14 days. U87-MG Glioblastoma Spheroids (a), SH-SY5Y Neuroblastoma spheroids (b), U87-MG + SH-SY5Y co-cultured spheroids (c)

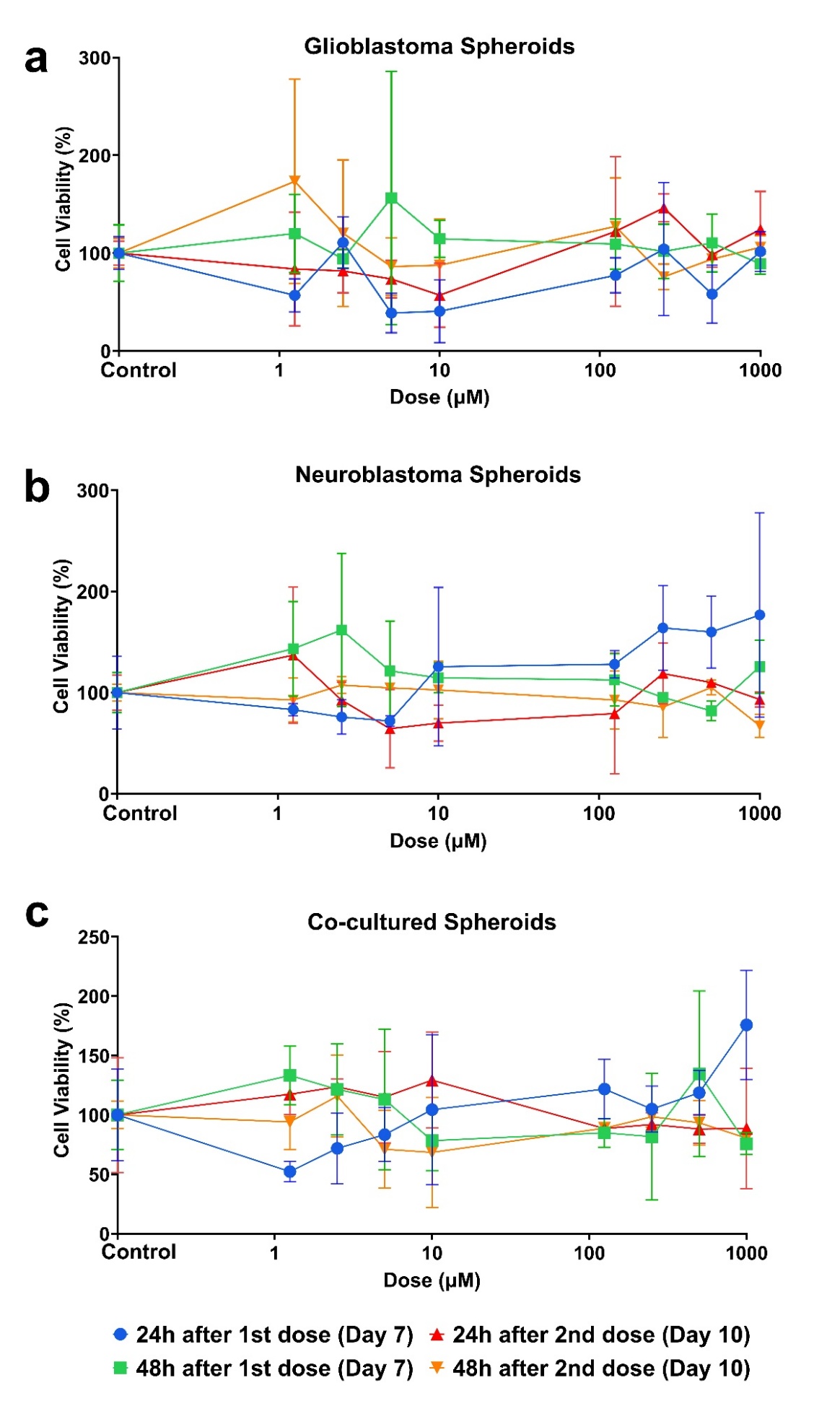

**Fig S3:** MTT Assay performed on spheroids after drug treatment with TMZ. U87-MG Glioblastoma Spheroids (a), SH-SY5Y Neuroblastoma spheroids (b), U87-MG + SH-SY5Y co-cultured spheroids (c)

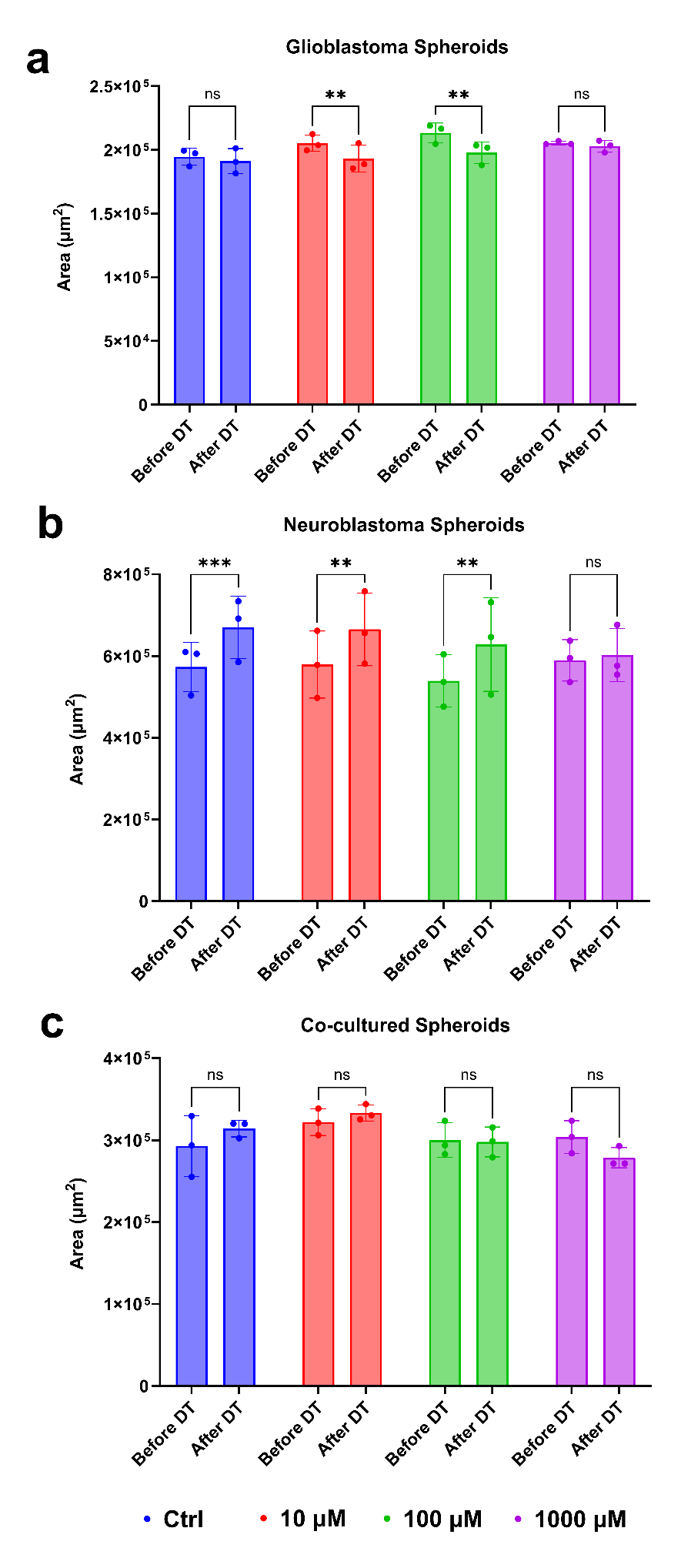

**Fig S4:** Change in area measured from the phase contrast images obtained before and after drug treatment on spheroids. U87-MG Glioblastoma Spheroids (a), SH-SY5Y Neuroblastoma spheroids (b), U87-MG + SH-SY5Y co-cultured spheroids (c)

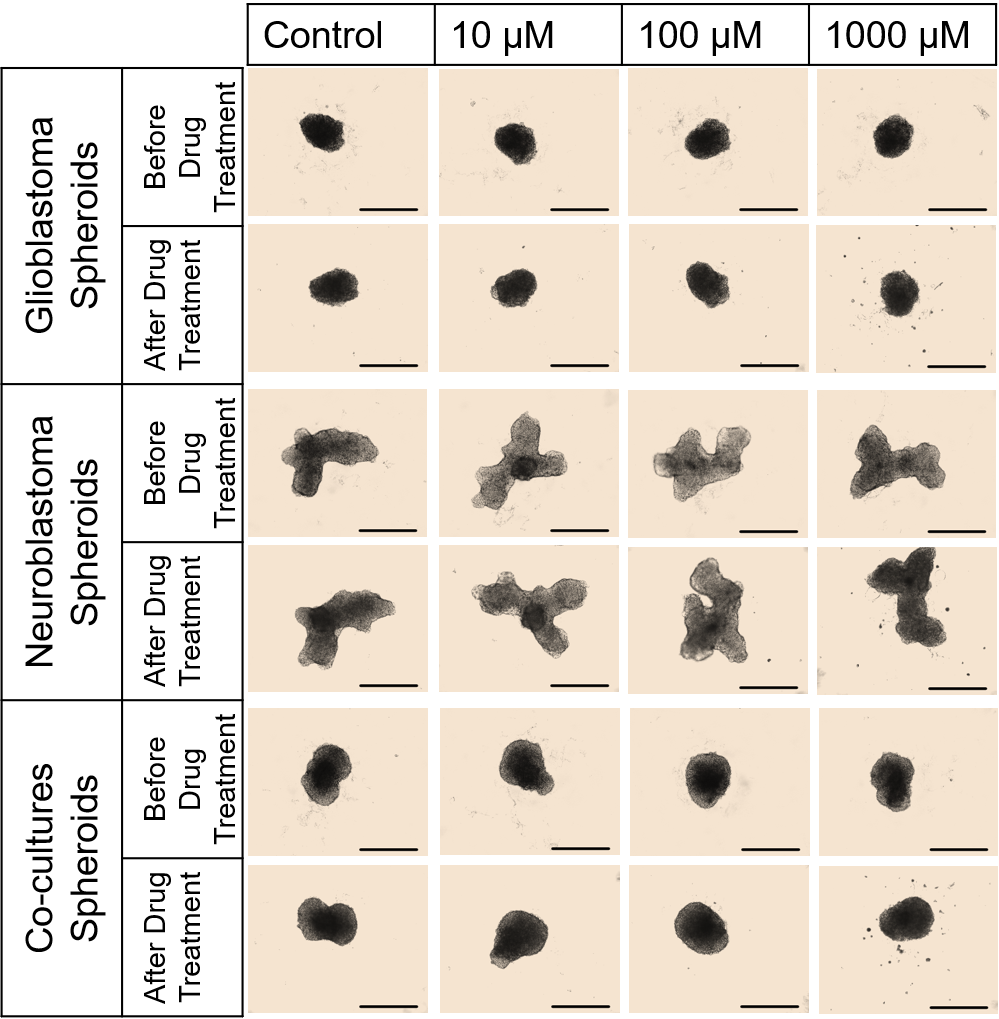

**Fig S5:** The phase contrast images obtained before and after drug treatment on spheroids. U87-MG Glioblastoma Spheroids (a), SH-SY5Y Neuroblastoma spheroids (b), U87-MG + SH-SY5Y co-cultured spheroids (c).Scale 500 μm
